# Dual PD-L1/TIGIT blockade induces PNAd⁺ HEV-like vessels and CD62L⁺ lymphocyte recruitment, driving rhabdoid tumor rejection

**DOI:** 10.64898/2025.12.15.692132

**Authors:** Stéphanie Fitte-Duval, Sofia Cavada Silva, Rafael Mena-Osuna, Owen Hoare, Leticia Laura Niborski, Jordan Denizeau, Laëtitia Lesage, Wilfrid Richer, Christine Sedlik, Kévin Beccaria, Maëva Veyssière, Rachida Bouarich-Bourimi, Jérémie Goldstein, Federico Marziali, Jimena Tosello Boari, Julien Masliah-Planchon, Zhi-Yan Han, Dario Rocha, Mylène Bohec, Sylvain Baulande, Joshua J. Waterfall, Valeria Manriquez, Franck Bourdeaut, Eliane Piaggio

## Abstract

Rhabdoid tumors (RTs) are aggressive pediatric malignancies with poor prognosis and limited immunotherapy options. Here, we investigate the therapeutic potential of combined PD-L1 (Programmed cell death ligand 1) and TIGIT (T cell immunoreceptor with Ig and ITIM domains) immune checkpoint blockade in RTs using a preclinical murine model that recapitulates key features of human ATRT (Atypical teratoid rhabdoid tumors) subtypes. Transcriptomic analyses of human and murine RTs reveal co-expression of TIGIT and PD-1 (Programmed cell death 1) pathway components and their ligands, particularly in immune-infiltrated subtypes, supporting a rationale for dual blockade. Combination therapy induces complete tumor regression, prolongs survival, and reprograms the tumor immune microenvironment by enriching CD62L⁺ naïve and central memory T cells and promoting selective T-cell clonal expansion. Notably, dual blockade initiates PNAd⁺ (Peripheral node addressin) high endothelial venule (HEV)-like structures, associated with focal lymphocyte clustering and enhanced immune cell recruitment. These findings reveal a mechanistic link between vascular remodeling and immune infiltration and support dual TIGIT and PD-L1 inhibition as a promising immunotherapeutic strategy for RTs.

## Introduction

Over recent decades, cancer immunotherapy has achieved remarkable successes^1^. Notably, immune checkpoint inhibitors (ICIs), including the inhibition of the PD-1/PD-L1 (Programmed cell death 1/ Programmed cell death ligand 1) pathway, have provided clinical benefits across multiple adult cancer types^2–4^.

The benefit of anti-PD1, anti-PD-L1 and anti-CTL4 treatments on pediatric cancers have been assessed within several pediatric trials^5^, with modest efficacy, with the exception of pediatric Hodgkin lymphoma^6^. This discrepancy may be attributed to differences in tumor immunogenicity, such as a lower mutational burden and reduced neoantigen load, as well as the relative immaturity of the immune system in children compared to adults^7,8^, which together may contribute to reduced responsiveness to immune checkpoint inhibitors. Nevertheless, occasional responses have been observed in pediatric patients with hypermutated or *SMARCB1*-deficient cancers^9–12^. Rhabdoid tumors (RTs), which are highly aggressive pediatric malignancies, are driven mainly by the biallelic inactivation of *SMARCB1*^13^. *SMARCB1* is a core component of the SWItch/Sucrose Non-Fermentable (SWI/SNF) complex, a major ATPase-dependent chromatin remodeler. RTs are classified as extracranial RTs (ECRTs) or extracranial malignant rhabdoid tumors (eMRTs) when arising outside the central nervous system (CNS), and as atypical teratoid/rhabdoid tumors (ATRTs) when arising within the CNS, where they comprise three epigenetically distinct subtypes: TYR, SHH and MYC^14,15^. Despite their genomic simplicity^16^, RTs can be infiltrated by immune cells^17–19^ and some individual cases have shown positive outcomes to ICIs, notably by PD-1/PD-L1 pathway inhibition^11,12,20^.

In a previously established mouse model of RT developed by our group^21^, PD-1/PD-L1 blockade significantly delayed tumor growth, but did not induce complete rejection in all cases^17^. Among promising combinatorial strategies to enhance therapeutic efficacy, co-targeting alternative immune checkpoints such as the TIGIT/CD226 axis has emerged as a promising approach^22^.

The TIGIT (T cell immunoreceptor with Ig and ITIM domains) receptor family (Fig. S1A), including TIGIT, CD226 (DNAM-1), CD96 (TACTILE), and PVRIG (CD112R), regulates T and NK (Natural killer) cell function with opposing effects: CD226 is co-stimulatory, while TIGIT, CD96, and PVRIG are co-inhibitory^23–25^. These receptors compete for shared ligands such as PVR (CD155), NECTIN2 (CD112) and NECTIN3 (CD113), expressed on tumor, stromal, and myeloid cells^22^. TIGIT suppresses anti-tumor immunity by impairing CD8⁺ T cell function, enhancing regulatory T cells (Treg) activity, and modulating macrophage responses^26–28^. It also synergizes with PD-1 and TIM3 to dampen T cell responses and limit anti-tumor efficacy^28,29^.

Therefore, co-blockade of TIGIT with either PD-1 or PD-L1 has been studied in murine cancer models, and shows tumor regression in B16F10 melanoma, MC38, CT26 colorectal carcinoma and GL261 glioblastoma mouse tumor models^29–33^. This effect has also been observed in pediatric neuroblastoma and medulloblastoma models^34,35^. Furthermore, several clinical trials investigating the combination of TIGIT and PD-1/PD-L1 blockade are currently underway in pediatric cancers including *SMARCB1*-deficient cancers (eg NCT05286801). Notably, among these trials, the CITYSCAPE study in non-small cell lung cancer (NSCLC) has demonstrated significant clinical responses, highlighting the therapeutic potential of combined TIGIT and PD-L1 blockade^36^.

While clinical trials have explored the combined blockade of PD-1/PD-L1 and TIGIT, its relevance in RTs remains unexplored. Here, we investigate the immunological mechanisms underlying this combined approach and demonstrate its efficacy in inducing complete tumor regression in a preclinical model of RTs.

## Results

### Transcriptomic analysis reveals expression patterns of TIGIT and PD-1 pathway members in human RTs

To characterize the expression patterns of ligands and receptors from the PD-1 and TIGIT immune checkpoint families in human RTs, we analyzed a transcriptomic dataset including bulk and single-cell RNA sequencing across multiple RT subtypes. The cohort includes 89 bulk RNA-seq samples and 6 single-cell RNA-seq data (scRNAseq) from tumor tissues, sorted immune populations, and longitudinal patient samples (Fig. 1A and Supplementary Data 1).

**Fig. 1.**
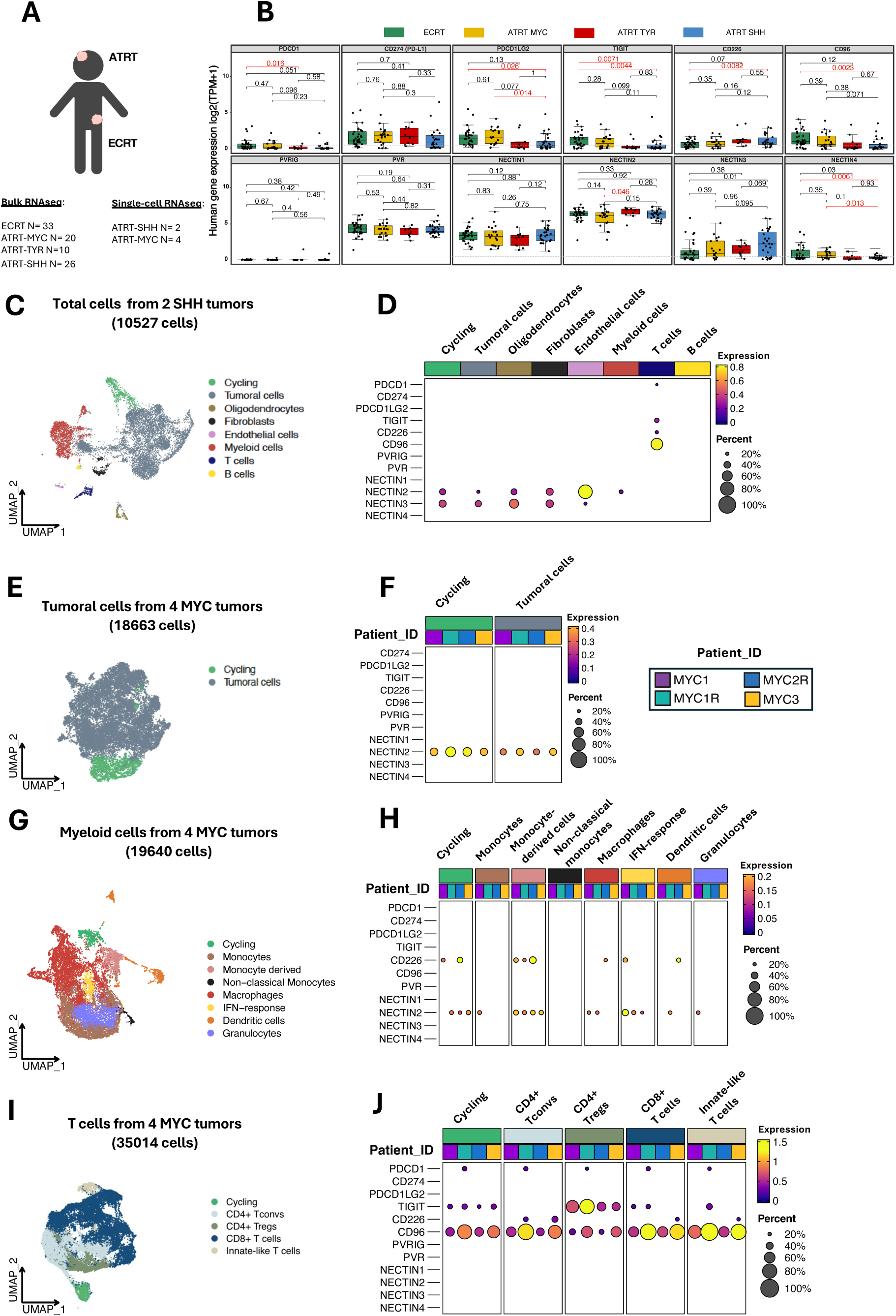
Transcriptomic analysis reveals expression patterns of TIGIT and PD-1 pathway members in human RTs. (A) Overview of analyzed samples. (B) Boxplots showing bulk RNA expression levels of genes related to the PD-1 and TIGIT family members obtained from human RT subtypes (ECRT N= 27, ATRT-MYC N= 20, ATRT-TYR N= 10, ATRT-SHH N= 25). Y axis shows expression as log2(TPM+1). Boxes represent the interquartile range (IQR), and whiskers extend to the smallest and largest values within 1.5 times the IQR below the first quartile (Q1) and above the third quartile (Q3), respectively. Significance was calculated using two-tailed Mann-Whitney test, with significant results (p < 0.05) indicated in red. (C) UMAP (uniform manifold approximation and projection) visualization of major cell types from integrated total cells of 2 tumors from patient SHH1 (10527 cells). (D) Single-cell RNA expression levels of PD-1/PDCD1 and TIGIT family genes across cell types from tumor microenvironment of patient SHH1. (E) UMAP visualization of integrated enriched tumoral cells from tumors of patients MYC1, MYC1R, MYC2R, MYC3 (18663 cells). (F) Single-cell RNA expression levels of *PD-1/PDCD1* and *TIGIT* family genes across tumor cells from 4 ATRT-MYC tumors. (G) UMAP visualization of integrated sorted myeloid cells from tumors of patients MYC1, MYC1R, MYC2R, MYC3 (19640 cells). (H) Single-cell RNA expression levels of *PD-1/PDCD1* and *TIGIT* family genes across sorted myeloid cells from 4 ATRT-MYC tumors. (I) UMAP visualization of integrated sorted T cells from tumors of patients MYC1, MYC1R, MYC2R, MYC3 (35014 cells). (J) Single-cell RNA expression levels of *PD-1/PDCD1* and *TIGIT* family genes across sorted T cells from 4 ATRT-MYC tumors. (D, F, H, J) Heatmaps showing SCT-normalized expression per gene and cell type. Dot size indicates the proportion of cells expressing the gene of interest within each cell type, and color indicates the level of expression from low (violet) to high (yellow). First row indicates the different cell types, using the same colors as in the corresponding UMAPs. Second row, labeled “Patient_ID”, uses colors to distinguish between different patients. Only genes expressed in more than 20% of cells are shown.

Bulk RNA-seq analysis revealed heterogeneous expression of ligands and receptors from the PD-1 and TIGIT families across RT subtypes (Fig. 1B and Supplementary Data 3). Among all RT samples, *CD274* (PD-L1), *PVR* (CD155), *NECTIN1* (CD111), *NECTIN2* (CD112), and *NECTIN3* (CD113) were the most highly expressed genes, whereas *PVRIG* was the only gene with undetectable expression. RTs generally exhibited low *PDCD1* (PD-1) expression but higher expression of its ligand *CD274*, supporting the rationale for therapeutic PD-L1 blockade. A trend toward higher expression of *PDCD1LG2* (PD-L2), *TIGIT*, *CD96*, and *NECTIN4* (PVRL4) was observed in ECRT and ATRT-MYC, previously identified as the most immune-infiltrated RT subtypes^17,18,37^, compared to ATRT-TYR and ATRT-SHH. Notably, immune receptors *PDCD1*, *TIGIT*, and *CD96* also showed elevated expression in these more infiltrated subtypes, whereas *CD226* (DNAM-1) exhibited the opposite pattern.

CD226 is a co-stimulatory receptor on lymphocytes that promotes effector function upon engagement with its ligands PVR and NECTIN2. These ligands, however, bind TIGIT with higher affinity, leading to inhibition of TIGIT^+^ T cell function. Among TIGIT co-receptors, *PVRIG*, an inhibitory receptor whose primary ligand is NECTIN2, was not detected. Conversely, *CD96*, which binds PVR and NECTIN1, showed a trend toward higher expression in ECRT and ATRT-MYC, mirroring the pattern observed for *TIGIT*, and could similarly contribute to immunosuppressive signaling. *CD226* was expressed at lower levels in ECRT and ATRT-MYC, and an inverse correlation was observed between *CD226* expression and *TIGIT*/*PDCD1*, likely reflecting the inhibitory effect of TIGIT and PD-1 on CD226 activation, which is critical for T/NK cell effector functions.

Overall, these bulk RNA-seq data indicate that RTs, particularly the more immune-infiltrated subtypes, express *Cd274* and *TIGIT* along with their main ligands *PVR* and *NECTIN2*. These expression patterns provide a mechanistic rationale for exploring combined PD-L1 and TIGIT blockade in RTs, with some subtypes potentially exhibiting a greater degree of pathway engagement than others.

While bulk RNA-seq captures gene expression from all cell types in the tumor microenvironment, it lacks cellular resolution. To address this, we utilized scRNAseq profiles to identify the cellular sources of PD-1/TIGIT pathway expression. We first analyzed scRNAseq data from the whole tumor microenvironment from two samples at two different progression disease timepoints obtained from one ATRT-SHH patient (SHH1). Eight major cell populations were identified, including Cycling cells (*MKI67⁺*), Tumoral cells (*SMARCB1⁻*, *SOX2⁺*), Oligodendrocytes (*OLIG1⁺*), Fibroblasts (*COL1A1⁺*), Endothelial cells (*PECAM1⁺*), Myeloid cells (*PTPRC⁺*; *CSF1R⁺* for monocytes/macrophages, *CSF3R⁺* for granulocytes, *FLT3⁺* for dendritic cells), T cells (*PTPRC*⁺, *CD3E*⁺), and B cells (*PTPRC*⁺, *CD19*⁺) (Fig 1C, S1B-C). Cell-type annotations were validated using established transcriptional signatures (Fig. S1D, Supplementary Data 2). Notably, although ATRT-SHH tumors showed overall lower expression of several PD-1/TIGIT pathway components in bulk RNAseq, scRNAseq reveal that some markers were still detectable in specific cell populations (Fig 1D). Consistent with the low or reduced expression in SHH bulk data, *CD274* and *PDCD1LG2* expression were undetectable in scRNAseq, whereas *PDCD1* was found only in a small proportion of T cells. *TIGIT* and its co-receptors *CD226* and *CD96*, but not their ligands, were present in T cells. Among TIGIT ligands, *NECTIN3* was restricted to tumoral and stromal cells (including oligodendrocytes, fibroblasts, and endothelial cells), whereas *NECTIN2* exhibited broader stromal expression, with notable enrichment in endothelial cells, and presence in myeloid cells.

To gain higher resolution of PD-1/TIGIT family expression per cell type, we next interrogated scRNAseq data from enriched tumor, sorted myeloid and T cell compartments from four ATRT-MYC samples to define the cellular sources of pathway activity. Analysis of enriched tumor cells revealed two major populations (Fig. 1E, S1E–F): Cycling (*MKI67⁺*) and Tumoral cells (*SMARCB1⁻ MYC⁺*) Only *NECTIN2* was consistently detected across tumor samples (Fig. 1F). No PD-1/PD-L1 family members were detected at the single-cell level in tumor cells, though *Cd274* was upregulated in some cases at the bulk RNA-seq level (Fig. 1B), likely reflecting expression from non-tumor cells within the tumor microenvironment included in the bulk data. These findings indicate that TIGIT ligands are expressed by ATRT-MYC tumor cells, while *Cd274* expression is likely contributed by the surrounding microenvironment.

scRNAseq analysis of sorted myeloid cells from the same four ATRT-MYC tumors previously analyzed, revealed eight major populations (Fig. 1G, S1G-H): Cycling (*MKI67^+^*), Monocytes (*CD14^+^ VCAN^+^*), Monocyte-derived cells (*CD14^-^ VCAN^-^HLA-DRA^+^*), Non-classical monocytes (*FCGR3A^+^*), Macrophages (*APOE^+^ CD14^+^ HLA-DRA^+^*), Interferon (IFN)-response (*RSAD2^+^*), Dendritic cells (*FLT3^+^*), and Granulocytes (*CSF3R^+^*). Cell-type annotations were validated using established transcriptional signatures (Fig. S1I, Supplementary Data 2). Among TIGIT and PD-1 ligands, only *CD226* and *NECTIN2* were expressed at low levels (Fig. 1H), suggesting that myeloid cells are unlikely to be primary target of the dual co-blockade in these tumors.

scRNAseq analysis of sorted T cells from the same four ATRT-MYC tumors previously analyzed, revealed five major T cell populations (Fig. 1I, S1J-K): Cycling (*MKI67^+^*), CD4^+^ conventional T cells (CD4^+^ Tconvs; CD3E^+^ CD4^+^ FOXP3^-^), CD4^+^ regulatory T cells (CD4^+^ Tregs; CD3E^+^ CD4^+^ FOXP3^+^), CD8^+^ T cells (*CD3E^+^ CD8A^+^*) and Innate-like T cells (*CD3E^+^ ZBTB16^+^ CD4^-^ CD8A^-^*). Cell-type annotations were validated using established transcriptional signatures (Fig. S1L, Supplementary Data 2). *PDCD1* was detected at more than 20% across T-cell subsets of only one sample, whereas *CD274* and *PDCD1LG2* were not expressed (Fig. 1J). *TIGIT* was present in all subsets, except CD4^+^ T convs, and was strongest in Tregs, while *CD226* remained consistently low. Notably, *CD96* was highly expressed across subsets, while *PVRIG* was not detected. Of note, CD96, with its dual inhibitory and activating motifs, can modulate immune responses context-dependently, potentially dampening activation by competing with CD226 for PVR binding ^38^. Lastly, TIGIT ligands (*PVR* and *NECTIN1–4*) were absent from T cells, indicating that ligand supply is extrinsic to T cells and suggesting a TIGIT/CD96-skewed checkpoint balance with limited CD226 co-stimulation.

Overall, our single-cell data suggest that anti-PD-L1 therapy may not benefit all RT patients, highlighting the need to stratify by PD-L1 expression. In contrast, TIGIT blockade could represent a broader strategy, given the widespread expression of its ligands NECTIN2 in myeloid and tumor cells and NECTIN3 in stromal and tumor cells, and consistent high TIGIT expression in Tregs.

In summary, expression patterns of TIGIT and PD-1 pathway members in human RTs support dual PD-L1 and TIGIT blockade as a promising therapeutic approach for immune-infiltrated RTs, particularly in ECRT and ATRT-MYC subtypes, and selected cases of other subtypes, warranting preclinical validation.

### Transcriptomic profile of the TIGIT/PD-1 axis in the RT mouse model supports the pre-clinical evaluation of dual immune checkpoint blockade

To evaluate the pre-clinical therapeutic efficacy of the dual blockade of TIGIT and PD-1, we used the RT mouse model developed by our group^21^, which recapitulates key features of human ATRT, including the anatomic and molecular characteristics of ATRT-MYC and ATRT-SHH subgroups *in situ*, as well as tumor immune infiltration in a syngeneic setting^17^. Additionally, blockade of the PD-1/PD-L1 pathway shows partial efficacy in this model^17^. First, we studied the expression pattern of TIGIT and PD-1 pathway members in the mouse tumor microenvironment (TME), following a similar approach as in humans. We analyzed a transcriptomic dataset including bulk and single-cell RNA sequencing of our available mouse RT subtypes, namely *in situ* and syngeneic ATRT-MYC and *in situ* ATRT-SHH (Fig. 2A, Supplementary Data 4). All ligands and receptors from the PD-1 and TIGIT families, except *Pvrig*, were detected in the mouse RT samples, with *Pdcd1*, *Cd274*, *Tigit*, *Cd226*, *Cd96*, *Pvr*, *Nectin2*, *Nectin4* being significantly more expressed in ATRT-MYC than in ATRT-SHH models (Fig. 2B).

**Fig. 2.**
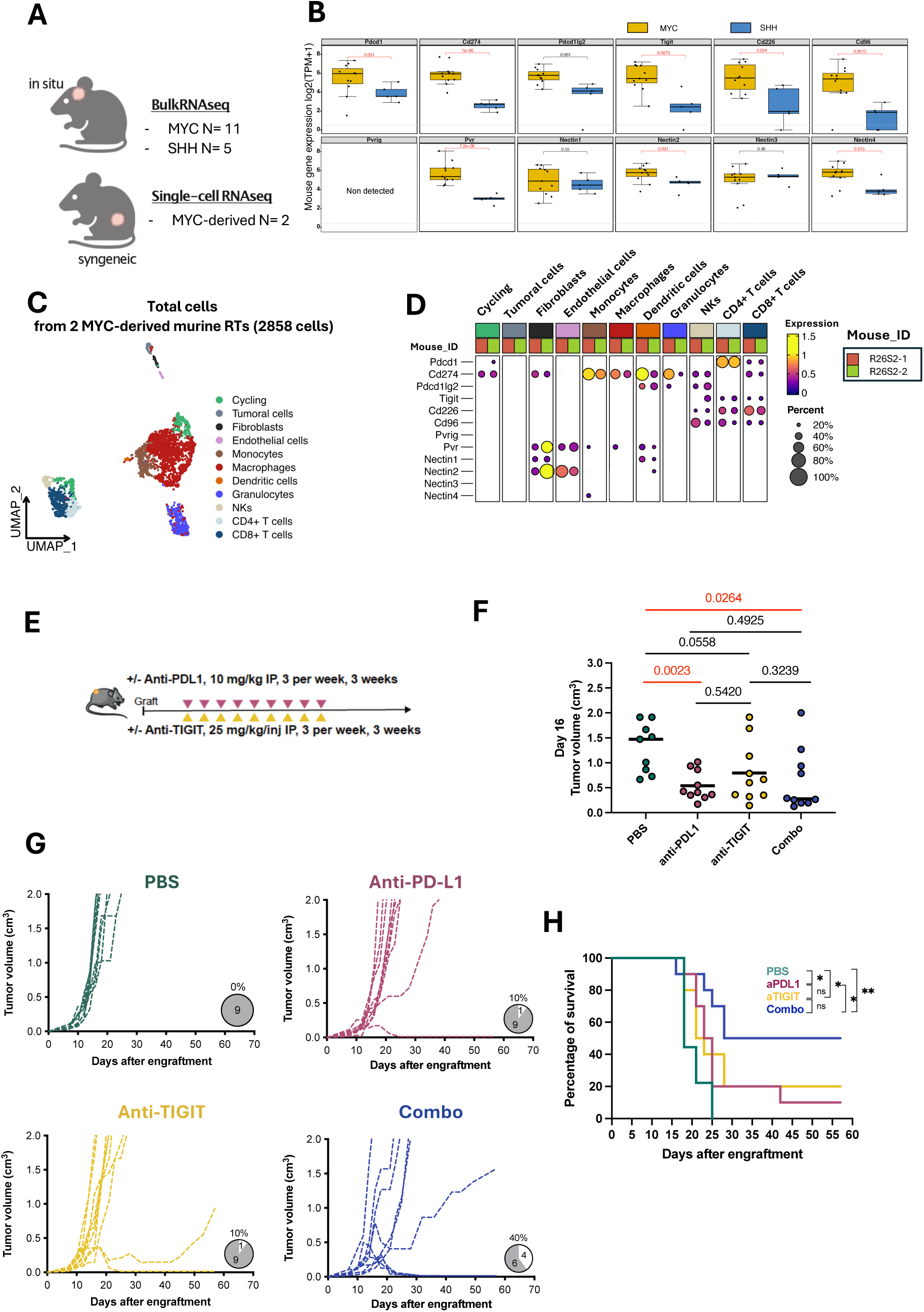
Transcriptomic profiling of the TIGIT/PD-1 axis in the RT mouse model supports its therapeutic targeting via dual immune checkpoint blockade. (A) Overview of murine samples used for transcriptomic analyses. (B) Boxplots showing bulk RNA expression levels of genes related to the PD-1 and TIGIT family members in the RT mouse model, according to molecular subgroups (ATRT-MYC, N= 11; ATRT-SHH, N= 5). Y axis shows expression as log2(TPM+1). Boxes represent the interquartile range (IQR), and whiskers extend to the smallest and largest values within 1.5 times the IQR below the first quartile (Q1) and above the third quartile (Q3), respectively. Significance was calculated using two-tailed Mann-Whitney test, with significant results (p < 0.05) indicated in red. (C) UMAP visualization of major cell types identified from integrated scRNAseq data (N= 2 syngeneic RT mice; total cells=2858). (D) Single-cell RNA expression levels of *PD-1/PDCD1* and *TIGIT* family genes cell types from tumor microenvironment from two syngeneic R26S2 RT mice. Heatmap shows SCT-normalized expression per gene and cell. Dot size indicates the proportion of cells expressing the gene of interest within each cell type, and color indicates the level of expression from low (violet) to high (yellow). First row indicates the different cell types, using the same colors as in the corresponding UMAPs. Second row, labeled “Mouse_ID”, uses colors to distinguish between different mice. Only genes expressed in more than 20% of cells are shown. (E) *In vivo* study design evaluating TIGIT and PD-L1 blockade. Mice were treated when tumor reached 2-4mm^2^ with intraperitoneal (IP) injections of anti-TIGIT or/and anti-PD-L1 at the indicated doses. (F-H) Results are representative of 2 independent experiments with 9-10 mice per group. (F) Tumor volume at day 16 (timepoint of first mouse sacrifice). Bars indicate median tumor volume per group. Statistical significance was calculated using two-tailed Mann-Whitney test, with significant results indicated in red. (G) Individual tumor growth curves for Control (PBS, N=9), anti-PD-L1 (N=10), anti-TIGIT (N=10), and combination (N=10). Pie charts indicate complete responses (white) and non-responses (grey), with numbers and percentages shown. (H) Kaplan-Meier survival curves. Statistical significance was calculated using the log-rank (Mantel-Cox) test (* = p < 0.05, ** = p < 0.01).

Additionally, we analyzed scRNAseq data from the whole TME of two Myc R26S2 RT samples. Eleven major cell populations were identified (Fig. 2C, S2A-B), including Cycling cells (*Mki67⁺*), Tumoral cells (*Smarcb1^-^ Ptprc^-^ Col1a1^-^ Pecam^-^),* Fibroblasts (*Col1a1^+^*), Endothelial cells (*Pecam1^+^*), Macrophages (*Ptprc^+^ Csf1r^+^ Fcgr1^+^*), Monocytes (*Ptprc^+^ Ly6c2^+^*), Granulocytes (*Ptprc^+^ S100a9^+^*), Dendritic cells (*Ptprc^+^ Flt3^+^*), NK cells (*Ptprc^+^ Ncr1^+^*), CD4^+^ T cells (*Ptprc^+^ Cd3e^+^ Cd4^+^*) and CD8^+^ T cells (*Ptprc^+^ Cd3e^+^ Cd8a^+^*). Cell-type annotations were validated using established transcriptional signatures (Fig. S2C, Supplementary Data 2).

*Pdcd1* expression was primarily restricted to T cells, with higher levels in CD4⁺ T cells (Fig. 2D). *Cd274* was expressed by fibroblasts and all immune populations, most strongly in myeloid cells, while *Pdcd1lg2* was found in dendritic cells, NK cells, T cells, and granulocytes. Tumor cells did not express PD-1 ligands. *Tigit* and its co-receptors *Cd226* and *Cd96* were detected in CD4⁺ and CD8⁺ T as well as NK cells, while *Pvrig* was not detected. Their ligands *Pvr* and *Nectin2* were mainly found in stromal cells and a subset of dendritic cells, *Nectin1* was detected in fibroblasts and dendritic cells, *Nectin4* showed low expression in monocytes, and *Nectin3* was not detected. Overall, the MYC-driven murine RT model shares key aspects of ATRT-MYC human data, including *Pdcd1*, *Tigit*, *Cd226*, and *Cd96* expression in T cells. While mice show higher Pvr and PD-1 ligands expression, taking this caveat into consideration, the model remains highly relevant for preclinical co-blockade studies targeting the PD-1/TIGIT axis.

### TIGIT/PD-L1 dual blockade enhances tumor rejection and survival in RT mouse model

To investigate the therapeutic efficacy of anti-PD-L1 and anti-TIGIT combination, RT-bearing mice were treated for three weeks with anti-PD-L1, anti-TIGIT, or their combination (Fig. 2E). Anti-PD-L1 treatment, both as monotherapy and in combination with TIGIT blockade, significantly delayed tumor growth by day 16 following treatment initiation, as evidenced by a marked reduction in tumor volume compared to control mice (Fig. 2F-G). Moreover, the combination therapy resulted in a higher proportion of mice achieving complete tumor rejection (Fig. 2G) which translated into prolonged overall survival relative to mice receiving monotherapies or control treatment (Fig. 2H). Taken together, these findings suggest a combined effect of anti-TIGIT and anti-PD-L1 therapies in enhancing anti-tumor responses in the mouse RT model.

### TIGIT/PD-L1 dual blockade reprograms tumor-infiltrating immune cells toward memory-like and functionally activated states

To assess how TIGIT/PD-L1 dual blockade shapes the tumor immune landscape, we profiled immune cells by scRNAseq and flow cytometry (Fig. 3A). Integration of 55,056 tumor-infiltrating CD45⁺ cells from 12 mice (N=3 per treatment group) identified seven major immune populations (Fig. 3B and S3A-B) including Cycling myeloid cells (*Mki67^+^ Ptprc^+^ Csf1r^+^ Fcgr1^+^ Flt3^-^*), Monocytes-Macrophages or MoMa, (*Ptprc^+^ Csf1r^+^ Fcgr1^+^ Flt3^-^*), Granulocytes (*Ptprc^+^ S100a9^+^*), Dendritic cells or DCs (*Ptprc^+^ Flt3^+^*), Cycling lymphoid cells (*Mki67^+^ Ptprc^+^ Cd3^+^ Trbc1^+^*), T/Natural Killer cells (T/NK cells) (*Ptprc^+^ Ncr1^+^ Cd3e^+^ Trbc1^+^*), and B cells (*Ptprc^+^ Cd19^+^ Flt3^-^*). Across treatments, relative abundance of these populations remained largely stable, except for a modest increase in intratumoral B cells (Fig. 3C-D, S3C-D). Thus, therapy did not substantially alter immune cell composition.

**Fig. 3.**
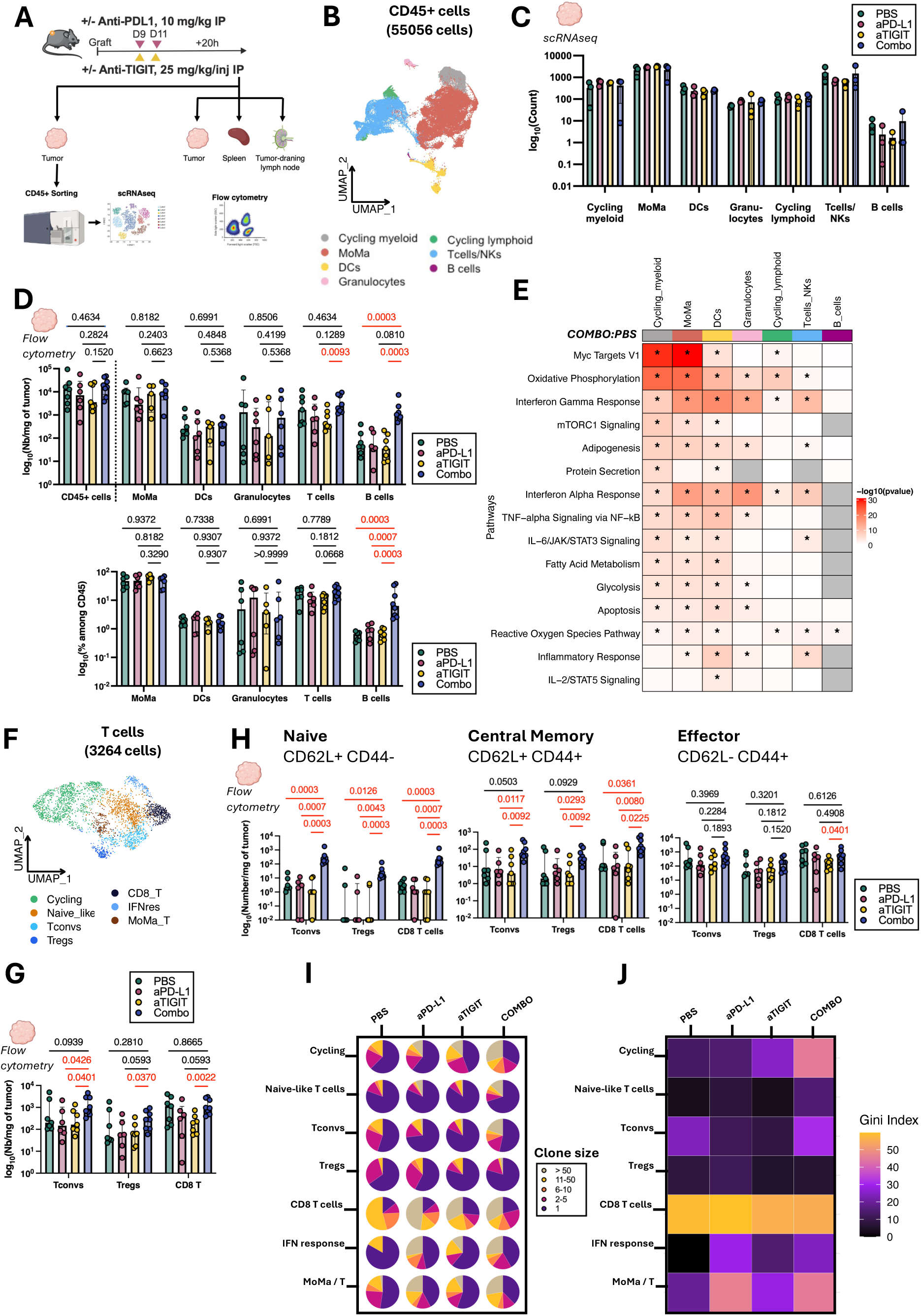
TIGIT/PD-L1 Dual Blockade Reprograms Tumor-Infiltrating Immune Cells Toward Memory-Like and Functionally Activated States. (A) *In vivo* study design evaluating the impact of TIGIT/PD-L1 blockade on immune composition. Tumor-bearing mice received two intraperitoneal injections of anti-TIGIT, anti-PD-L1, or both, and tumors were collected 20 hours after the final injection. (B-C; E-F; I-J) Single-cell RNA sequencing analysis of RT mouse tumors treated with PBS, anti-TIGIT, anti-PD-L1, or combined therapy (N=3 per treatment group). (D; H-G) Flow cytometry analysis of RT mouse tumors treated with the different regimens. Lymphoid panel: PBS (N=7), anti-PD-L1 (N=6), anti-TIGIT (N=7), and combination therapy (N=8). Myeloid panel: PBS (N=6), anti-PD-L1 (N=6), anti-TIGIT (N=5), and combination therapy (N=6). Bars represent median values; error bars indicate the interquartile range (IQR). (B) UMAP visualization of major immune cell types identified from integrated CD45^+^ sorted cells (N=3 per treatment group; total cells=55056). (C) Quantification of the major immune cell populations across treatment groups from single-cell RNA sequencing. Bars represent mean values; error bars indicate the standard deviation (SD). (D) Quantification (number/mg of tumor) and frequency (%) of major immune cells isolated from murine tumors by flow cytometry. (E) Pathway-enrichment analysis in major immune cell populations comparing Combo-and PBS-treated tumors. (F; I-J) Single-cell RNA sequencing analysis of T cells isolated from B). (G-H) Flow cytometry analysis of T cell subpopulations (Tconvs, Tregs, CD8^+^ T cells) isolated from murine tumors. (F) UMAP visualization (N= 3 per treatment group; total cells=3264) (G) Quantification (number/mg of tumor) of T cell subpopulations by flow cytometry. (H) Quantification (number/mg of tumor) by flow cytometry of Naïve (CD62L^+^ CD44^-^), Central Memory (CD62L^+^ CD44^+^) and Effector Memory (CD62L^-^ CD44^+^) Tconvs, Tregs and CD8^+^ T cells. Statistical significance was calculated using two-tailed Mann-Whitney test. Significant results indicated in red in flow cytometry barplots and single-cell violin-plots and with a star (p-value < 0.05) in pathway-enrichment heatmaps. (I) Pie charts showing relative expansion of TCR clonotypes among T cell subpopulations across treatments. Slices represent clone sizes ranging from 0 to > 50. (J) Heatmap of Gini index of TCR clonotype distributions within T cell subpopulations across treatment. Higher Gini values (warmer colors) indicate reduced clonal diversity, reflecting a more oligoclonal repertoire driven by dominant clonotypes. Rows represent T cell subsets and columns the treatment groups.

We next examined cell-intrinsic functional states, focusing on Combo versus PBS (Fig. 3E, Supplementary Data 5). Dual blockade induced broad metabolic and inflammatory rewiring, including enrichment of Myc and mTORC1 signaling in cycling cells, MoMa and DCs, activation of IFN responses, TNF-α/NF-κB and IL-6/JAK/STAT3 signaling, and increased mitochondrial ROS (Reactive oxygen species) across immune populations, including B cells. In contrast, T cells showed limited proliferative expansion but extensive metabolic and transcriptional reprogramming. These data indicate that therapy primarily reprograms immune function rather than abundance.

To identify the cellular sources of these inflammatory programs, we next analyzed MoMa, DC, and T cells at the subpopulation level. The abundance of the identified MoMa subpopulations remained largely stable (Fig. S3E-G) while they exhibited functional plasticity (Fig. S3H). Specifically, upon dual blockade, cycling, IFN responding (IFNres) and macrophages upregulated proliferative pathways and most subpopulations showed strong IFN- and NF-κB-driven responses. In addition, cycling and IFNres displayed secretory activity and regulatory signatures (*Tgfb1*, *Spp1*; Supplementary Data 6), suggesting the presence of simultaneous pro-inflammatory and homeostatic activity.

Similarly, the different identified DC subpopulations did not significantly change in abundance (Fig. S3I-K) but were broadly reprogrammed toward pro-inflammatory and antigen-presenting states (*Il12b*, *Il18*, *H2-K1*; Supplementary Data 6) (Fig. S3L). Specifically, following anti-TIGIT plus anti-PD-L1 treatment, cDC1 and cDC2 (conventional DC 1 & 2) subsets upregulated antigen processing and co-stimulatory pathways *(Tap1*, *Cd40, Cd86*; Supplementary Data 6), while MoDCs (Monocyte-derived Dendritic cells) and DC3s (Dendritic cells 3) induced chemokine programs *(Ccl3, Cxcl9*; Supplementary Data 6) potentially facilitating T cell recruitment.

For T cells, we identified both classical subsets (Cycling, Tconvs, Tregs, CD8⁺) and functional populations not strictly defined by CD4⁺/CD8⁺ identity, including naïve-like, IFN-responsive (IFNres), and MoMa/T hybrids (Fig. 3F, S3M-N). Stable abundance across most subsets was observed, with a tendency to an increase in intratumoral Tconvs under dual treatment (Fig. 3G).

At the functional level, dual blockade remodeled metabolic and inflammatory pathways across T cell subsets (Fig. S3O). Cycling, IFNres, and MoMa/T subsets were the most activated, with enrichment of OXPHOS (Oxydative Phosphorylation), Myc Targets v1, interferon, and inflammatory pathways (IL-2/STAT5, TNFα/NF-κB; Fig. S3O). CD8⁺ T cells and Tconvs also showed metabolic activation, together with central memory–associated markers (*Klf2*, *Sell*, *Ccr7*, *Lef1*, *Ly6c2*^39^ (Fig. S3P, Supplementary Data 6), indicating preferential commitment to memory-like rather than terminal effector states. Tregs remained modestly activated and naïve-like T cells largely quiescent. Together, these data suggest that dual blockade predominantly redirects T cells toward central memory programs rather than driving full effector differentiation, potentially supporting durable antitumoral immunity despite restrained proliferative expansion.

To confirm the transcriptional changes, we assessed the distribution of naïve (CD62L⁺CD44⁻), central memory (CD62L⁺CD44⁺), and effector/effector memory (CD62L⁻CD44⁺) subsets within Tconvs, Tregs, and CD8⁺ T cells by flow cytometry. Combination treatment led to increased infiltration of Tconvs, Tregs, and CD8⁺ T cells with a naïve or central memory phenotype, while effector subsets remained comparable to controls (Fig. 3H). This shift resulted in a significant higher proportion of naïve and central memory T cells at the expense of effector cells (Fig. S3Q), corroborating the single-cell findings of restrained effector expansion and enhanced memory bias.

Finally, we evaluated clonal expansion. Consistent with upregulation of the “Myc Targets v1” pathway, indicative of T cell proliferation, most T cell subsets expanded after single or dual checkpoint blockade, except naïve-like T cells and Tregs (Fig. 3I). Gini index analysis, which quantifies clonotype distribution inequality^40^, revealed that dual blockade led to greater clonal dominance than controls, indicating a shift toward fewer, more expanded clones. Overall, dual blockade enhances the antitumor T cell response by driving the expansion of dominant clones (Fig. 3J).

Immune composition in spleen and lymph nodes was largely preserved across treatments, with only modest increases in Ly6C⁺ monocytes, and monocyte-derived cells upon dual blockade (Fig. S3R-T).

Overall, these findings indicate that TIGIT/PD-L1 dual blockade reprograms tumor-infiltrating immune cells without substantially altering their overall subsets composition. Within tumors, myeloid and dendritic cells adopt pro-inflammatory, antigen-presenting states, while T cells undergo metabolic and transcriptional remodeling with selective expansion of dominant clones. This shift favors naïve and central memory phenotypes, producing a more clonal, memory-biased antitumor T cell response.

### Combination blockade of PD-L1 and TIGIT induces the formation of PNAd⁺ HEV-like structures in the tumor microenvironment

High endothelial venules (HEVs), characterized by the expression of peripheral node addressin (PNAd), mediate lymphocyte recruitment into lymphoid tissues via interactions with L-selectin (CD62L)^41,42^, which is expressed on naïve T cells, central memory T cells, and some B cells - all of which were increased following combination therapy (Fig. 3H, S3Q). In tumors, HEV-like vessels are often associated with the formation of tertiary lymphoid structures (TLS)-organized aggregates of B cells, T cells, and follicular dendritic cells, frequently localized around PNAd⁺ HEVs. The presence of TLS and HEV-like vasculature has been linked to improved immune cell infiltration and enhanced responses to immunotherapy^43–45^. Based on this, we investigated whether dual PD-L1 and TIGIT blockade could promote lymphoid neogenesis in the tumor microenvironment. Immunofluorescence staining for PNAd revealed very limited HEV-like structures in tumors from control or monotherapy-treated mice. In contrast, tumors from mice receiving the combination treatment exhibited a notable, though variable, increase in PNAd⁺ structures, with several displaying vascular morphology (Fig. 4A-B). Although this increase did not reach statistical significance, the trend suggests that dual checkpoint blockade may favor the initiation of HEV-like remodeling. In control and monotherapy groups, T cells and CD11c⁺ dendritic cells were diffusely distributed across the tumor parenchyma, which showed minimal PNAd expression (Fig. 4A-B). In the combination treated group, however, we observed focal clusters of immune cells adjacent to PNAd⁺ vessels, with a tendency toward increased local densities of B and CD11c⁺ cells. Of note, a significant increase in the abundance and frequency of CD62L⁺ B and T cells (Fig. 4C, S4) was observed only in the combination-treated group, correlating with the higher number of PNAd⁺ cells and consistent with the enrichment of B cells, as well as naïve and central memory lymphocytes in tumors. Moreover, CD3⁺ TCF1⁺ T cells and CD11c⁺ cells were often found within 20 μm of PNAd⁺ structures (Fig. 4D), suggesting a spatial association between emerging vascular structures and recruitment of immune cells.

**Fig. 4.**
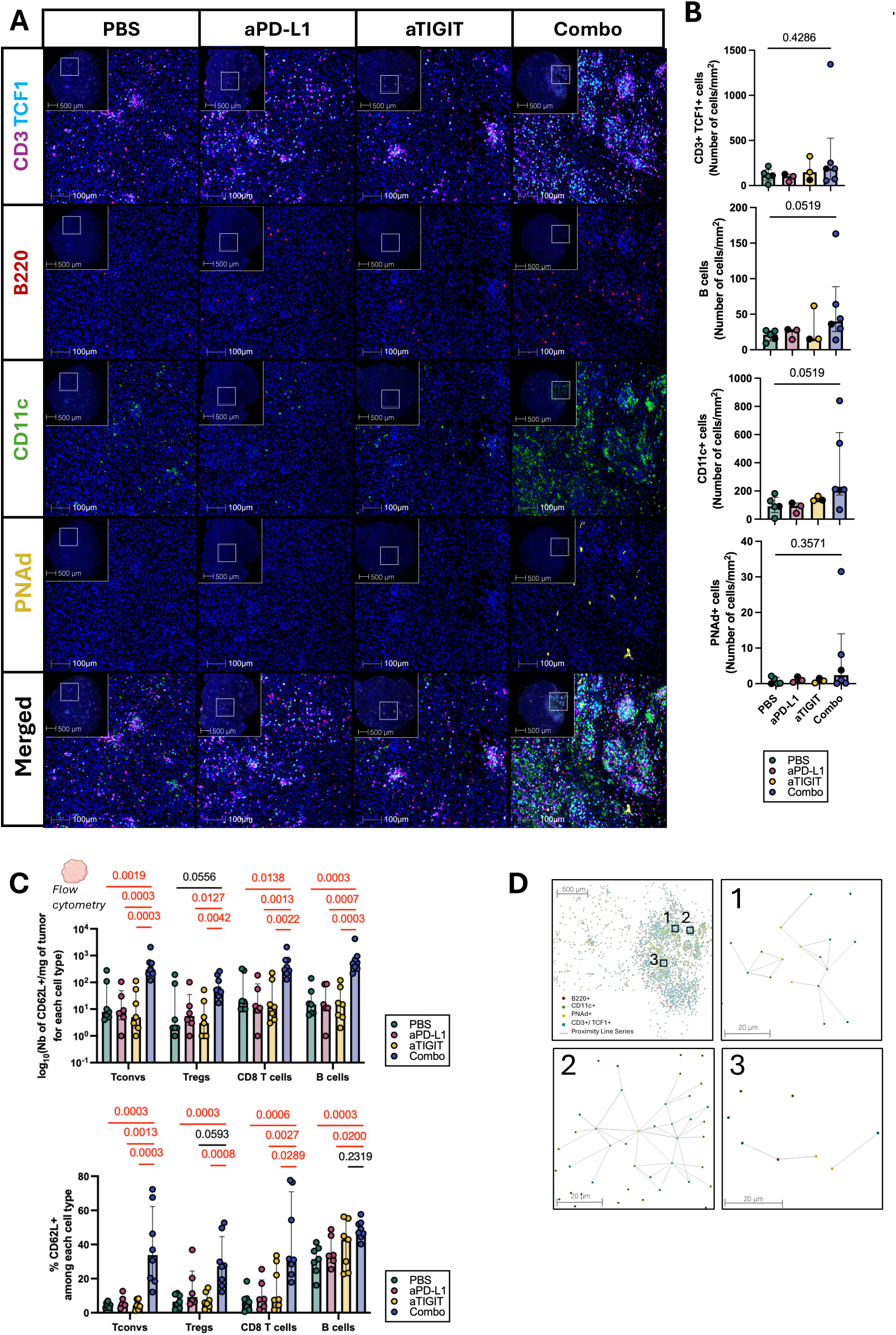
Combination therapy promotes HEV-like remodeling. (A) Multiplex immunohistochemistry (mIHC) staining of a representative tumor per treatment condition showing the co-expression of CD3 (magenta), TCF1 (cyan), B220 (red), CD11c (green), PNAd (yellow) with nuclei counterstained with DAPI (blue). White scale bars=1 mm. (B) Quantification of cell densities per treatment condition. PBS (N=5), anti-PD-L1 (N=3), anti-TIGIT (N=3), and combination therapy (N=6). Black dots indicate the displayed sample per condition in (A). Bars represent the median cell count per group; error bars indicate the interquartile range (IQR). Statistical significance was calculated using two-tailed Mann-Whitney test, with significant results indicated in red. (C) Quantification (number/mg of tumor) and frequency (%) of CD62L^+^ B and T cells from immune cells isolated from murine tumors by flow cytometry across treatment: PBS (N=7), anti-PD-L1 (N=6), anti-TIGIT (N=7), and combination therapy (N=8). Bars represent the median cell count per group; error bars indicate the interquartile range (IQR). Statistical significance was calculated using two-tailed Mann-Whitney test, with significant results indicated in red. (D) Representative composite images depicting proximity analysis between PNAd^+^ cells (yellow) and CD3^+^ TCF1^+^ cells (blue), CD11c^+^ cells (green) and B cells (pink). The proximity lines show proximity of CD3⁺TCF1⁺, CD11c⁺, and B cells to each other (<20 µm) in a radius under 20 µm to PNAd⁺ cells.

Interestingly, tumors in the combination group appeared to fall into two phenotypic patterns: those with increased PNAd expression, higher density of B and CD11c⁺ cells, and higher proportion of T cells expressing CD62L; and those resembling the low-infiltrate profile seen in controls. This heterogeneity may reflect variation in the extent of lymphoid neogenesis across tumors and could potentially relate to the variability in therapeutic response previously observed (Fig. 2G–H).

Overall, despite the emergence of immune-rich niches, we did not detect mature TLS, which are typically organized lymph-node-like structures containing B-cell follicles, T-cell zones, DCs, and specialized vasculature^46^. This suggests that TLS development may be at an early stage or that the subcutaneous tumor model may not fully support TLS maturation, as previously reported^43,45^.

## Discussion

Our study demonstrates that dual PD-L1 and TIGIT blockade induces complete tumor regression in a preclinical model of rhabdoid tumors (RTs), a rare pediatric malignancy with few effective therapeutic options. We show that this combination therapy remodels the tumor immune landscape by increasing the recruitment of naïve and central memory T cells, while promoting selective T cell clonal expansion and the emergence of PNAd⁺ high endothelial venule (HEV)-like structures, suggesting that these vascular and immunological changes may underlie the observed curative responses.

Transcriptomic analyses across human RT subtypes revealed co-expression of TIGIT and its ligands (NECTIN2, NECTIN3) across immune and tumor compartments, and to a lesser extent PD-L1, particularly in immune-infiltrated tumors such as ATRT-MYC and ECRT subgroups. This aligns with reports that TIGIT and PD-1 are frequently co-expressed on dysfunctional tumor-infiltrating lymphocytes (TILs) in adult cancers and can cooperatively suppress effector functions through CD226 antagonism^26,36,47^. Importantly, our preclinical RT model recapitulates this immunogenomic landscape, validating its utility for testing TIGIT-based combinatorial approaches.

Consistent with prior work in colorectal cancer^28,48^, HNSCC models^49^, glioblastoma^33^, and neuroblastoma^34^, our data show that TIGIT/PD-L1 dual blockade synergistically promotes tumor rejection. Mechanistically, this was associated with increased infiltration of CD4⁺ and CD8⁺ T cells harboring naïve and central memory phenotypes, leading to a therapeutic benefit driven by recruitment and expansion of naïve-like, memory-precursor, and effector cells as previously demonstrated with ICIs^50,51^.

A key insight from our study is the induction of PNAd⁺ HEV-like vessels in the combined treated group, which facilitate lymphocyte entry into secondary lymphoid organs via L-selectin/CD62L interactions^52^. The increased presence of CD62L⁺ T and B cells in treated tumors supports active lymphocyte trafficking and correlates with improved tumor control^53^. HEVs and the broader process of lymphoid neogenesis have been increasingly recognized as favorable features in immuno-oncology, correlating with response to ICIs in melanoma, NSCLC, and breast cancer^42,54–57^. While we observed immune cell clustering near HEV-like vessels, the absence of mature tertiary lymphoid structures (TLS) suggests an early stage of lymphoid organization or possible limitations of the subcutaneous tumor model for supporting full TLS maturation^43,57,58^.

Our findings align with a growing body of literature highlighting the role of vascular remodeling and stromal activation in shaping tumor immunity^45,59,60^. Recent work has shown that immune checkpoint blockade can trigger vascular remodeling and TLS formation, enhancing antigen presentation and T cell priming^61,62^. Notably, the spatial proximity of TCF1⁺ stem-like T cells and CD11c⁺ dendritic cells to PNAd⁺ structures in our model suggests a nascent lymphoid niche conducive to local T cell activation, a hallmark of TLS biology.

The synergistic effects of TIGIT and PD-L1 blockade likely result from their convergent inhibition of co-stimulatory signaling. Both pathways impair CD226 activation, a central axis for cytotoxic T cell and NK cell function, via ligand competition (NECTIN2/3) and phosphatase recruitment^63^. By releasing this inhibition, dual blockade may restore CD226-dependent activation and promote the generation of functional memory T cells, as shown in other models^22,50,64^. Moreover, TIGIT signaling in dendritic cells has been implicated in suppressing IL-12 production and antigen presentation, effects that may also be reversed by anti-TIGIT therapy^30^.

Importantly, not all tumors responded uniformly to treatment; some displayed limited HEV induction and immune infiltration. This intragroup heterogeneity mirrors observations in clinical studies and underscores the need to identify biomarkers predictive of response. Future work should determine whether PNAd⁺ vessel induction or the presence of naïve/central memory T cells can serve as early correlates of efficacy. Furthermore, given the restricted maturation of TLS in our model, alternative tumor settings (e.g., orthotopic or spontaneous models) may be required to evaluate the full potential of lymphoid neogenesis as a therapeutic endpoint.

In summary, our data support dual PD-L1 and TIGIT blockade as a rational and potent immunotherapeutic strategy for RTs. By facilitating the recruitment and reprogramming of lymphocytes, and promoting vascular niches reminiscent of lymphoid structures, this approach offers a path to long-term tumor control and immune memory in an otherwise refractory tumor type. These findings provide a preclinical foundation for advancing TIGIT-based combinations in pediatric solid tumors and highlight HEV induction and immune reorganization as promising avenues for therapeutic exploitation.

## Methods

### Human Tumor Samples

Freshly resected and snap-frozen human RT samples were collected following written informed consent of parents regarding tumor banking and use for research; approval of these consents was obtained by with approved protocol N°2021-A00795-36 from Committee for the Protection of Individuals (“Comité de protection des personnes”). All samples were pseudo-anonymized prior to processing.

### Mice

C57BL/6J female mice were obtained from Charles River Laboratories, maintained in a non-barrier facility and included at 8-12 weeks of age for experimental procedures. Spontaneous tumors were obtained using the conditional Smarcb1^flox/flox^; Rosa26-Cre^ERT2^ system as previously described ^21^, where biallelic Smarcb1 deletion is obtained upon tamoxifen injection at E6-E7.

Syngeneic R26S2 mouse RT models were derived from the spontaneous model and were maintained by successive subcutaneous engraftments in the flank of C57BL/6J mice, every 4-5 weeks. Animal care and use for this study were performed in accordance with the recommendations of the European Community (2010/63/UE) for the care and use of laboratory animals. Experimental procedures were specifically approved by the ethics committee of the Institut Curie CEEA-IC #118 (Authorization APAFiS#47128-2024012511115844-v2 given by National Authority) in compliance with the international guidelines.

### Mice Treatments

For treatment experiments using the syngeneic RT models, 10-20 mm^3^ piece of tumor per mice were engrafted subcutaneously in the flank. 7 days after engraftment, mice with measurable tumor size between 2-4 mm^2^ were included in the experiment. Groups were stratified by tumor size to have equivalent mean size between the groups on the first day of treatment. RT-bearing mice were treated with anti-PD-L1 (10mg/kg, clone 6E11, Roche Genentech), anti-TIGIT (25mg/kg, clone 10A7, Roche Genentech), or a combination of both three times a week, for three weeks. Control mice received PBS. Tumor volume was determined by caliper measurements using the formula: V = a × b^2^/2 with a being the largest diameter and b the smallest. Survival was assessed using the time to reach a humane endpoint, defined as a tumor volume of 2 cm³ or a body weight loss exceeding 10%.

For flow cytometry, single-cell and immunohistochemistry experiment upon treatment, 2 doses of anti-PD-L1 and anti-TIGIT, or combination of both were administered at days 9 and 11 post-engraftment. After 20 hours, tumors were collected for further flow cytometry, single-cell, or immunohistochemistry analysis, and spleens and tumor-draining lymph nodes (tdLNs) for flow cytometry only.

### Mice tissue dissociation

Tumors (for flow cytometry and single-cell analysis), spleens and tdLNs (for flow cytometry) were harvested. Tumors and spleens were digested in Digestion mix, containing 2 mL of CO₂-independent medium (Gibco) supplemented with 0.1 mg/mL DNase I (Roche) and 0.1 mg/mL Liberase™ TL (Roche) at 37 °C, using a gentleMACS™ Dissociator (Miltenyi Biotec) with the pre-set program “37C_m_TDK1” (41 min at 37 °C under continuous agitation). Collected lymph nodes were mechanically dissociated prior to enzymatic digestion for 45 min at 37 °C in Digestion mix. Enzymatic reactions were stopped by adding MACS buffer (0.5% BSA, bovine serum albumin), and 2 mM EDTA (Invitrogen) in PBS 1X (Eurobio-Scientific), and cell suspensions were filtered through a Falcon® 100 µm cell strainer. Red blood cells were removed by treatment with RBC lysis buffer (2 min in 2 ml from 1L of purified water supplemented with 8,32g of NH_4_Cl, 0,84g of NaHCo_3_, 0,043g of EDTA, from Sigma-Aldrich). The resulting single-cell suspensions were washed, resuspended in MACS buffer, and the number of viable cells was determined using acridine orange staining with an automated cell counter (LUNA-FX7, Logos Biosystems).

### Flow cytometry analysis

Live/dead cell discrimination was performed using Live/Dead Fixable Aqua Dead Cell Stain Kit (Life Technologies) and Fc receptors were blocked with the anti-mouse CD16/CD32 (clone 2.4.G2) mAb (BD). When possible, samples were split in half to stain with either a Lymphocyte or a Myeloid Panel, using the antibodies listed in Table S1. Cell surface labelling was performed at 4°C and intracellular staining was performed after using Fixation/Permeabilization Kit (eBioscience/Thermo Fischer) according to the manufacturer’s instructions. Data acquisition was performed using an LSR Fortessa cytometer (BD) and analyzed using FlowJo software 10.10.0 (TreeStar). GraphPad Prism version 10.2.1 was used for data visualization.

### Single-Cell RNA sequencing

#### Human samples

First sample of live cells from SHH1 patient, FACS-enriched CD45^-^ cells and myeloid cells (CD45^+^ CD3^-^) from patient MYC1, MYC1R, MYC2R and MYC3 were loaded on a Chromium X (10X Genomics) and libraries were prepared using a Single Cell 3′ Reagent Kit V3 (10X Genomics) according to the manufacturer’s protocol. Second sample of live cells from SHH1 patient and FACS-enriched T cells (CD45^+^ CD3^+^ CD19^-^) from patient MYC1, MYC1R, MYC2R and MYC3 were loaded on a Chromium X (10X Genomics) and libraries were prepared using a Single Cell 5′ Reagent Kit V2 (10X Genomics) according to the manufacturer’s protocol.

#### Mouse samples from co-blockade experiment

FACS-enriched CD45^+^ cells from 12 treated mice were loaded on a 10X Chromium (10X Genomics) and libraries were prepared using a Single Cell 5′ Reagent Kit V2 (10X Genomics) according to the manufacturer’s protocol.

Only for FACS-enriched T cells from mouse samples, V(D)J segments were selectively enriched prior to library construction. Library quantification and quality assessment were achieved using a dsDNA High Sensitivity Assay Kit (Thermo Fisher Scientific) and a Bioanalyzer 2100 system (Agilent Technologies). 3’ or 5’ gene expression libraries were tested for quality, equimolarly pooled and sequenced on an Illumina NovaSeq platform using paired-end as sequencing mode. Enriched V(D)J libraries were sequenced on an Illumina MiSeq system.

### Single-cell data analysis

Samples used for the human and mouse scRNAseq cohorts are listed in Supplementary Data 1 ^17,21^. To obtain count matrices, human scRNAseq data were processed with Cellranger 7.1.0 using GRCh38, and mouse scRNAseq data were processed with Cellranger 6.0.0 using mm10 as genome reference (refdata-gex-mm10-2020-A). Recovery of neutrophils was performed using Cellranger count with the parameter --force-cells=40000. Resulting count matrices of each sample were first analyzed and integrated by Seurat v4, then the integrated objects and downstream figures were analyzed with Seurat v5 in R v4.4^65^. For each sample, all cells expressing more than 10% as mitochondrial gene percentage were considered as dead cells and were excluded from downstream analysis. Only for the two whole tumor samples of SHH1 that were low quality, all cells expressing more than log_10_(3) of total UMI counts were maintained for downstream analysis. For the CD45^+^ mouse cells from co-blockade experiment, neutrophils were first identified given their characteristically low transcript counts, to perform appropriate nFeature filtering. A minimum threshold of 1,000 detected features was then applied to all non-neutrophil cells. At this stage, clusters displaying a high proportion of mitochondrial transcripts, as well as cells with persistently low nFeature values, were also removed. For each sample, contaminant red blood cells were removed based on *HBB*, *HBA1*, *HBA2* genes.

Normalization was performed with Seurat’s SCTransform^66,67^. Data integration across samples used Seurat’s anchor-based workflow (FindIntegrationAnchors() and IntegrateData()) to harmonize gene expression without additional batch correction^68^.

Dimensionality reduction was achieved with PCA (RunPCA()), retaining the most informative components. For T cells containing datasets, TCR genes were excluded from the analysis to avoid biases from clonal expansion.

Graph-based clustering was performed with the Louvain algorithm (FindClusters()) and visualized via UMAP (RunUMAP()), with clustering stability assessed using Clustree v0.5.1.

For *Human enriched tumoral cells dataset (Fig. 1E)*, the following clusters were eliminated: Stromal cells (PECAM1), oligodendrocytes (OLIG1), Cancer-Associated Fibroblasts (FAP), immune cells (CD45).

For *Human sorted myeloid cells dataset (Fig. 1G)*, the following clusters were eliminated: Stromal cells (PECAM1, COL1A1), T cells (CD45, CD3)

For *Human sorted T cells dataset (Fig. 1I)*, the following clusters were eliminated: Stromal cells (PECAM1, COL1A1), Myeloid cells (CD45, CD3neg).

All downstream analyses, including normalization, integration, clustering, and visualization, were then repeated after contaminant removal.

### Cell type annotation

Cell clusters were annotated using FindAllMarkers() (Wilcoxon test, adj. P < 0.05, |log2FC| > 0.25) and canonical markers using FeaturePlot() and ViolinPlot(), with cluster identities further validated using published gene signatures and differential gene expression (Supplementary Datas S3-S4) quantified via AddModuleScore() and displayed by DotPlot().

### Bulk mRNA cohorts’ analysis

Samples used for the human mRNA-seq cohort are listed in Supplementary Data 1. Samples used for the mouse mRNA-seq cohort are listed in Supplementary Data 1 and were obtained from the Smarcb1^flox/flox^; Rosa26-Cre^ERT2^ model (Shh, n = 5; Myc, n = 11) described previously^21,69^. Bulk RNA-seq samples were classified according to tumor subtype (ECRT, ATRT-MYC, ATRT-TYR, ATRT-SHH for patient samples, and Myc or Shh for mouse samples).

RNA sequencing (RNA-seq) data were preprocessed and analyzed using the Institut Curie RNA-seq Nextflow pipeline (v4.1.0; https://doi.org/10.5281/zenodo.7443721). In brief, raw reads were trimmed with TrimGalore (v0.6.7), aligned to the hg38 human reference genome with STAR, and quantified with FeatureCounts (v1.4.0) using the Gencode v45 annotation. Violin plots of gene expression data were produced using the log2 transformation of transcripts per million (TPM) data + 1 with the R package ggplot2 (v.3.5.0) (Supplementary Datas S3-S4). Significance was assessed using the Mann-Whitney test implemented in R v4.4.

### Gene expression analysis of PD1 and TIGIT family members

Gene expression of PD1 and TIGIT family members was analyzed using the GroupHeatmap() function from the SCP package (v0.5.6, https://zhanghao-njmu.github.io/SCP/). To corroborate the visualization, the mean expression of each gene of interest was calculated across cells with expression >0, and the percentage of expressing cells was also determined for each cell subpopulation and patient in Supplementary Datas 3-4. As the expressions were similar for the two samples from SHH1, the samples were merged for this analysis.

### Downsampling and count analysis

For comparisons across samples from co-blockade experiment, all samples from the integrated dataset were down sampled to the number of cells in the smallest sample. Cells were randomly selected without replacement from each sample to match this minimum cell count, ensuring equal representation across conditions for downstream analyses. UMAP and GraphPad Prism version 10.2.1 was used for data visualization.

### Pathways Enrichment analysis

For each dataset, differentially expressed genes with a threshold of p-value < 0.05 and a |log₂FC| > 0.5 between Combo and PBS in each cell subset were identified using FindMarkers() and listed in Supplementary Datas S5-S6. Upregulated genes were subjected to pathway enrichment analysis via the Enrichr API, using the MSigDB database. Enrichment metrics of all terms, including overlap score, p-value, adjusted p-value, odds ratio, combined score, and associated genes were recorded in Supplementary Datas S5-S6. P-values were transformed using -log10 for heatmap visualization, with terms ranked according to the p-values of the first cell subset in each dataset. Statistical significance was assessed using the Mann–Whitney test, with significant results indicated by a star (p < 0.05).

### Subsets analysis

To further investigate immune cell subpopulations, two datasets were extracted from the integrated Cd45⁺ object based on annotations: (1) “Cycling myeloid”/“MoMa”, (2) “DCs”. To gain more T cells content, the T cell dataset has been obtained by identifying T cells (Cd3e^+^, Trac^+^, Trbc1^+^, Cd8a^+^ or Cd4^+^) in each sample and by integrating them. NK cells and innate-like T cells were not included in this analysis as no significant changes in single-cell or flow cytometry analyses were observed.

For each dataset, down sampling, count analysis, and pathway enrichment analysis were performed. In the T cell dataset, functional gene expression was further assessed, with average expression and the percentage of expressing cells visualized using DotPlot().

### Single-Cell T Cell Receptor Analysis

V(D)J sequences were assembled with Cell Ranger v7.1.0 (10x Genomics) using the refdata-cellranger-vdj-GRCm38-alts-ensembl-7.0.0 reference. Using R, clonotypes were defined by unique CDR3 nucleotide sequences and V subunits, retaining only full-length, productive TRA and TRB contigs; cells with >1 TRB or >2 TRA chains were excluded as potential doublets.

To increase the number of cells with TCR information (only 695 cells had TCR data in the down sampled dataset), cells removed during the down sampling process were recovered from the raw dataset containing both RNA and VDJ data. A total of 2,865 cells with RNA and TCR information absent from the down sampled dataset were identified. These cells were annotated using a label transfer approach, applying the FindTransferAnchors() function followed by TransferData(). After this procedure, downstream clonal analyses were performed on 3,560 cells. Clonal expansion was quantified by counting cells per clonotype (defined by TRB chain by combining CDR3 with V subunits) and analyzed across treatment conditions and T cell subpopulations. Clonal composition and relative abundance were visualized using pie charts. Clonal diversity was assessed using the Gini-TCR Skewing Index ^40^, calculated separately for each subpopulation and treatment group.

### Multiplexed immunohistochemistry

Tumors were collected in 4% paraformaldehyde (Edimex, 3178-200-19), incubated for 24h in the dark and transferred into 80% ethanol for slides preparation by the platform of Experimental pathology of Institut Curie. Alcohol, formalin, and acetic acid (AFA)-fixed paraffin-embedded tissue blocks were cut with a microtome into fine slivers of 3 microns. Immunostaining was processed in a Bond RX automated (Leica) with Opal™ 7-Color IHC Kits (Akoya Biosciences, NEL821001KT) according to the manufacturer’s instructions. The multiplex panel consisted of the following antibodies: CD11c (Cell Signaling, #97585, 1/200), CD3 (Agilent, GA5036, RTU), TCF1 (Cell signaling, #2203, 1/100), B220 (BD Biosciences, 553086, 1/400), CD31, (Abcam, ab28364, 1/100) and PNAd (BD Biosciences, 553863, 1/500). Tissue sections were coverslipped with Prolong™ Diamond Antifade Mountant (Thermo Fisher) and stored at 4°C. Subsequently, slides were scanned using the Vectra ® 3 automated quantitative pathology imaging system (Vectra 3.0.5; Akoya Biosciences). Multispectral images were unmixed using inForm advanced image analysis software (v2.6, Akoya Biosciences). Cell quantification and proximity analysis were performed using Halo software (v3.5; Indica Labs).

### Statistical Analysis

The tests used for statistical analyses are described in the legends of each concerned figure and have been performed using GraphPad Prism version 10.2.1 or R v4.4. For comparing the expression of PD-1, TIGIT and their ligands between the different RT subtypes in bulk analyses, tumor growth, flow cytometry and single-cell counts, p-values were calculated using the nonparametric Mann-Whitney test. Statistical significance was defined by p < 0.05 (two-tailed). Data are presented as medians ± IQR. For mouse survival studies, p-values were calculated using the log-rank (Mantel-Cox) test (* = p < 0.05, ** = p < 0.01).

### Data availability

All data supporting the findings of this study will be deposited in EGA and ArrayExpress and made publicly available upon publication. Data deposition is currently underway and will be accessible for peer review. Bulk RNA-seq data of mouse rhabdoid tumor model are accessible through GEO accession number GSE241734.

## Acknowledgements

This work was supported by Roche Genentech, PRTK-19-052 funding from INCa and DGOS, the LabEx DCBIOL (ANR-10-IDEX-0001-02 PSL; ANR-11-LABX-0043); and Center of Clinical Investigation (CIC IGR-Curie 1428). This work was carried out within the framework of Eliane Piaggio’s team labeled by the Fondation pour la Recherche Médicale (FRM, grant number EQU202203014661). We would like to thank SFCE-Enfants Cancer Santé, Marabout de Ficelles and SMARCB1 hope for supporting all our research for rhabdoid tumors. We also acknowledge La Ligue contre le cancer for funding the PhD programs of Stéphanie Fitte-Duval and Sofia Cavada Silva. High-throughput sequencing was performed by the ICGex NGS platform of the Institut Curie supported by the grants ANR-10-EQPX-03 (Equipex) and ANR-10-INBS-09-08 (France Génomique Consortium) from the Agence Nationale de la Recherche (“Investissements d’Avenir” program), by the ITMO-Cancer Aviesan (Plan Cancer III) and by the SiRIC-Curie program (SiRIC Grant INCa-DGOS-465 and INCa-DGOSInser12554). We would like also to acknowledge the Institut Curie’s CurieCoreTech platforms for their technical and scientific expertise, including the animal facility, cytometry, NGS and Experimental pathology platforms. We also acknowledge Raphaël Hartmann, Jérémy Mesple for technical support, Annaëlle Marey for administrative support, and Sergio Roman Roman and Pedro Cerda Hernandez for helpful discussions.

## Author contributions

Conceptualization and methodology: SFD, CS, VM, FB, EP

Anti-PD-L1 & Anti-TIGIT blockade material supply: Roche Genentech

Experiment: SFD, SCS, RMO, LLN, RBB, JTB, FM, JD, CS, VM

Clinical samples: KB

R26S2 mouse model generation & bulkRNAseq data collection: ZYH

Patient bulkRNAseq collection: JMP & FB

BulkRNAseq data analysis: OH, SFD

Single-cell RNAseq data analysis: SFD, SCC, OH, WR, DR, VM

Single-cell TCRseq data analysis: SFD, SCC, WR, VM

Data management and R-language support: MV, WR

NGS support: MB, SB

Histological data acquisition and analysis: LL, SFD

Supervision: JJW, VM, FB, EP

Writing of original draft: SFD, JJW, VM, FB, EP

Manuscript proofreading: CS

Funding acquisition: JG, VM, FB, EP

All authors reviewed and contributed to the final manuscript

## Competing interests

EP is co-founder of Egle Therapeutics. EP and JJW are share-holders at Mnemo Therapeutics. The remaining authors declare no competing interests.

**Correspondence** and requests for materials should be addressed to Valeria Manriquez, Franck Bourdeaut or Eliane Piaggio.

