## Supplementary Figure for "Dual PD-L1/TIGIT blockade induces PNAd⁺ HEV-like vessels and CD62L⁺ lymphocyte recruitment, driving rhabdoid tumor rejection"

**The PDF file includes:**

Supplementary Figures:

Fig. S1. Transcriptomic analysis reveals subtype-specific expression patterns of TIGIT and PD-1 pathway members in human RTs.

Fig. S2. Transcriptomic profile of the TIGIT/PD-1 axis in the RT mouse model supports the pre-clinical evaluation of dual immune checkpoint blockade.

Fig. S3. TIGIT/PD-L1 Dual Blockade Reprograms Tumor-Infiltrating Immune Cells Toward Memory-Like and Functionally Activated States.

Fig. S4. Combination blockade of PD-L1 and TIGIT induces the formation of PNAd+ HEV-like structures in the tumor microenvironment.

Supplementary Table:

Table S1. List of antibodies used in this study.

**Other Supplementary Material for this manuscript includes the following:**

Data Supplementary 1 - Sample information

Data Supplementary 2 - Signatures for single-cell annotations

Data Supplementary 3 - Human bulk and single-cell data - Related to Fig. 1

Data Supplementary 4 - Murine bulk and single-cell data - Related to Fig. 2

Data Supplementary 5 - scRNAseq of tumors from treated mice - Related to Fig.3

Data Supplementary 6 - scRNAseq of tumors from treated mice (continuation) - Related to Fig. S3

### Supplementary Figures

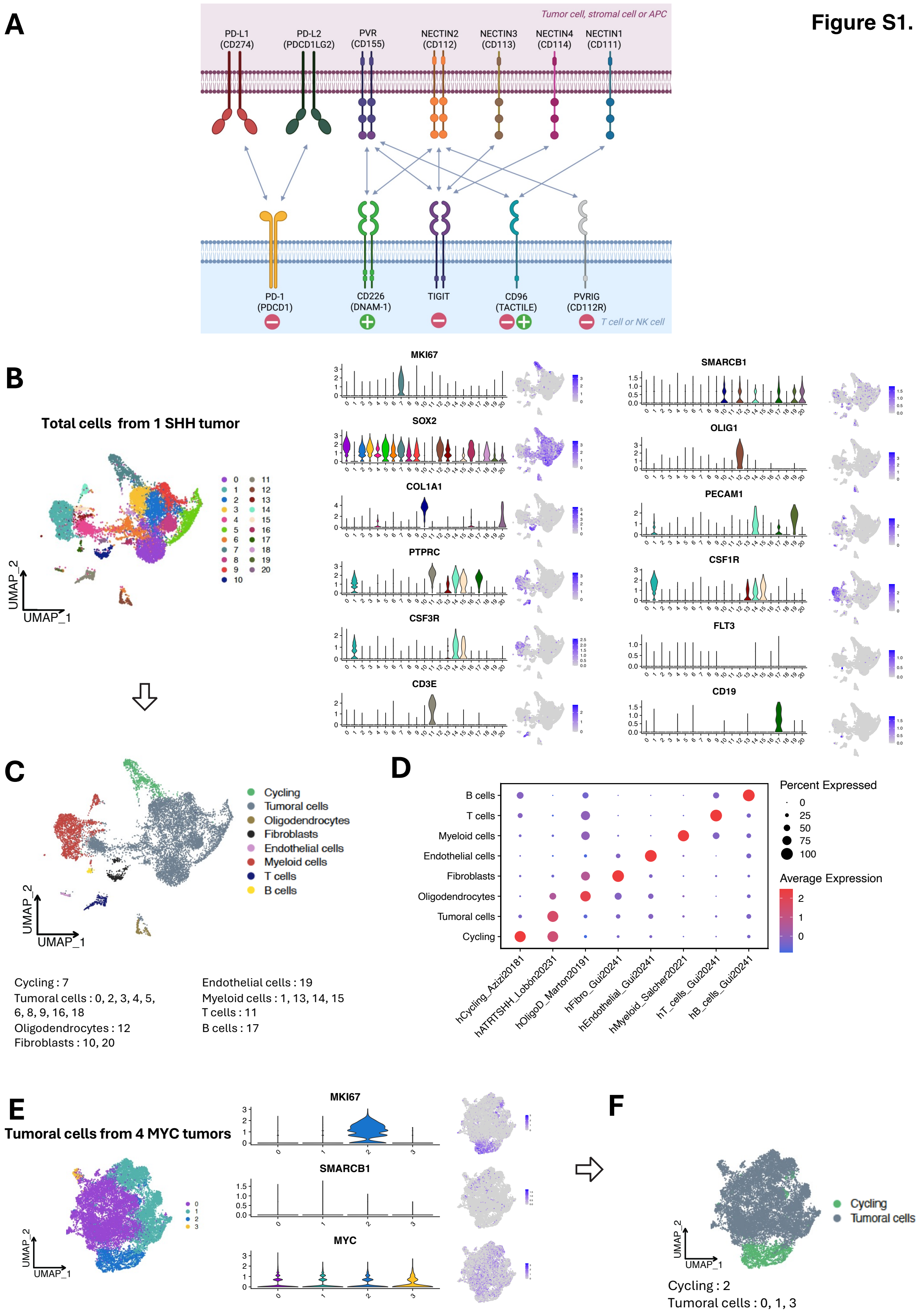


**Figure S1.**

**Fig. S1 (related to Fig. 1). Transcriptomic analysis reveals expression patterns of TIGIT and PD-1 pathway members in human RTs.**

(A) Scheme representing interactions of PD-1 and TIGIT receptors' family. Created with Biorender.

(B-D) Workflow for annotating cell populations from single-cell RNA-seq of total cells from tumors of patient SHH1.

(B) UMAP from integrated total cells from 2 tumors of patient SHH1 at resolution 0.7 before annotation (10527 cells). Violin and feature plots showing the expression pattern of representative genes across clusters.

(C) Merging of similar clusters into consolidated and annotated cell populations based on shared gene expression profiles.

Cluster 7 was renamed "Cycling". Clusters 0, 2, 3, 4, 5, 6, 8, 9, 16 and 18 were merged into "Tumoral cells". Cluster 12 was renamed "Oligodendrocytes". Clusters 10, 20 were merged into "Fibroblasts". Cluster 19 was renamed "Endothelial cells". Clusters 1, 13, 14 and 15 were merged into "Myeloid cells". Cluster 11 was renamed "T cells". Cluster 17 was renamed "B cells"

(D) Dot plot showing the average gene expression per annotated cluster to validate cluster annotation.

(E-F) Workflow for annotating cell populations from single-cell RNA-seq of enriched tumoral cells from 4 ATRT-MYC tumors.

(E) UMAP from integrated total cells from 4 ATRT-MYC tumors at resolution 0.1 before annotation (18663 cells). Violin and feature plots showing the expression pattern of representative genes across clusters.

(F) Merging of similar clusters into consolidated and annotated cell populations based on shared gene expression profiles.Cluster 2 was named "Cycling". Clusters 0, 1, 3 were merged into "Tumoral cells".

**Figure S1 (2).**

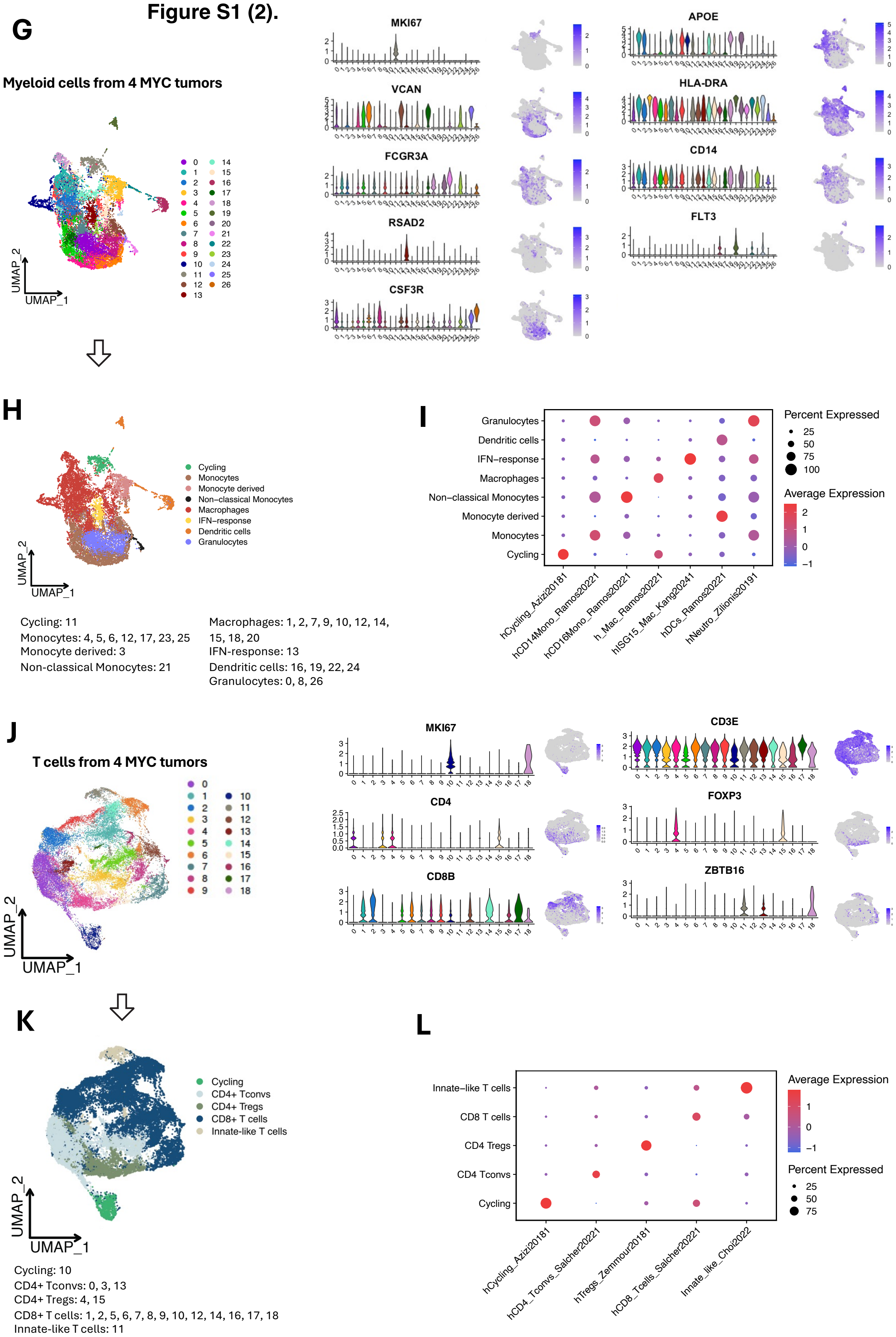
G
Myeloid cells from 4 MYC tumors
UMAP_2
UMAP_1
MKI67
VCAN
FCGR3A
RSAD2
CSF3R
APOE
HLA-DRA
CD14
FLT3
H
Cycling
Monocytes
Monocyte derived
Non-classical Monocytes
Macrophages
IFN-response
Dendritic cells
Granulocytes
Cycling: 11
Monocytes: 4, 5, 6, 12, 17, 23, 25
Monocyte derived: 3
Non-classical Monocytes: 21
Macrophages: 1, 2, 7, 9, 10, 12, 14, 15, 18, 20
IFN-response: 13
Dendritic cells: 16, 19, 22, 24
Granulocytes: 0, 8, 26
I
Percent Expressed
Average Expression
hCycling_Azizi20181
hCD14Mono_Ramos20221
hCD16Mono_Ramos20221
h_Mac_Ramos20221
hISG15_Mac_Kang20241
hDCs_Ramos20221
hNeutro_Zilionis20191
J
T cells from 4 MYC tumors
MKI67
CD3E
CD4
FOXP3
CD8B
ZBTB16
K
Cycling
CD4+ Tconvs
CD4+ Tregs
CD8+ T cells
Innate-like T cells
Cycling: 10
CD4+ Tconvs: 0, 3, 13
CD4+ Tregs: 4, 15
CD8+ T cells: 1, 2, 5, 6, 7, 8, 9, 10, 12, 14, 16, 17, 18
Innate-like T cells: 11
L
Innate-like T cells
CD8 T cells
CD4 Tregs
CD4 Tconvs
Cycling
Average Expression
Percent Expressed
hCycling_Azizi20181
hCD4_Tconvs_Salcher20221
hTregs_Zemmour20181
hCD8_Tcells_Salcher20221
Innate_like_Choi2022

**Continuation of Fig. S1 (related to Fig. 1). Transcriptomic analysis reveals expression patterns of TIGIT and PD-1 pathway members in human RTs**

(G-I) Workflow for annotating cell populations from single-cell RNA-seq of sorted myeloid cells from 4 ATRT-MYC tumors.

(G) UMAP from integrated sorted myeloid cells from 4 ATRT-MYC tumors at resolution 1 before annotation (19640 cells). Violin and feature plots showing distribution of representative genes for each cluster.

(H) Merging of similar clusters into consolidated and annotated cell populations based on shared gene expression profiles.

Cluster 11 was renamed "Cycling". Clusters 4, 5, 6, 12, 17, 23, 25 were merged into "Monocytes". Cluster 3 was renamed "Monocyte derived". Cluster 21 was renamed "Non-classical Monocytes". Clusters 1, 2, 7, 9, 10, 12, 14, 15, 18, 20 were merged into "Macrophages". Cluster 13 was renamed "IFN-response". Clusters 16, 19, 22, 24 were merged into "Dendritic cells". Clusters 0, 6, 26 were merged into "Granulocytes".

(I) Dot plot showing the average gene expression per annotated cluster to validate cluster annotation.

(J-L) Workflow for annotating cell populations from single-cell RNA-seq of sorted T cells from 4 ATRT-MYC tumors.

(J) UMAP from integrated sorted T cells from 4 ATRT-MYC tumors at resolution 0.5 before annotation (35014 cells). Violin and feature plots showing distribution of representative genes for each cluster.

(K) Merging of similar clusters into consolidated and annotated cell populations based on shared gene expression profiles.

Cluster 10 was renamed "Cycling". Clusters 0, 3, 13 were merged into "$CD4^+$ Tconvs". Clusters 4, 15 were merged into "$CD4^+$ Tregs". Clusters 1, 2, 5, 6, 7, 8, 9, 10, 12, 14, 16, 17, 18 were merged into "$CD8^+$ T cells". Cluster 11 was renamed "Innate-like T cells".

(L) Dot plot showing the average gene expression per annotated cluster to validate cluster annotation.

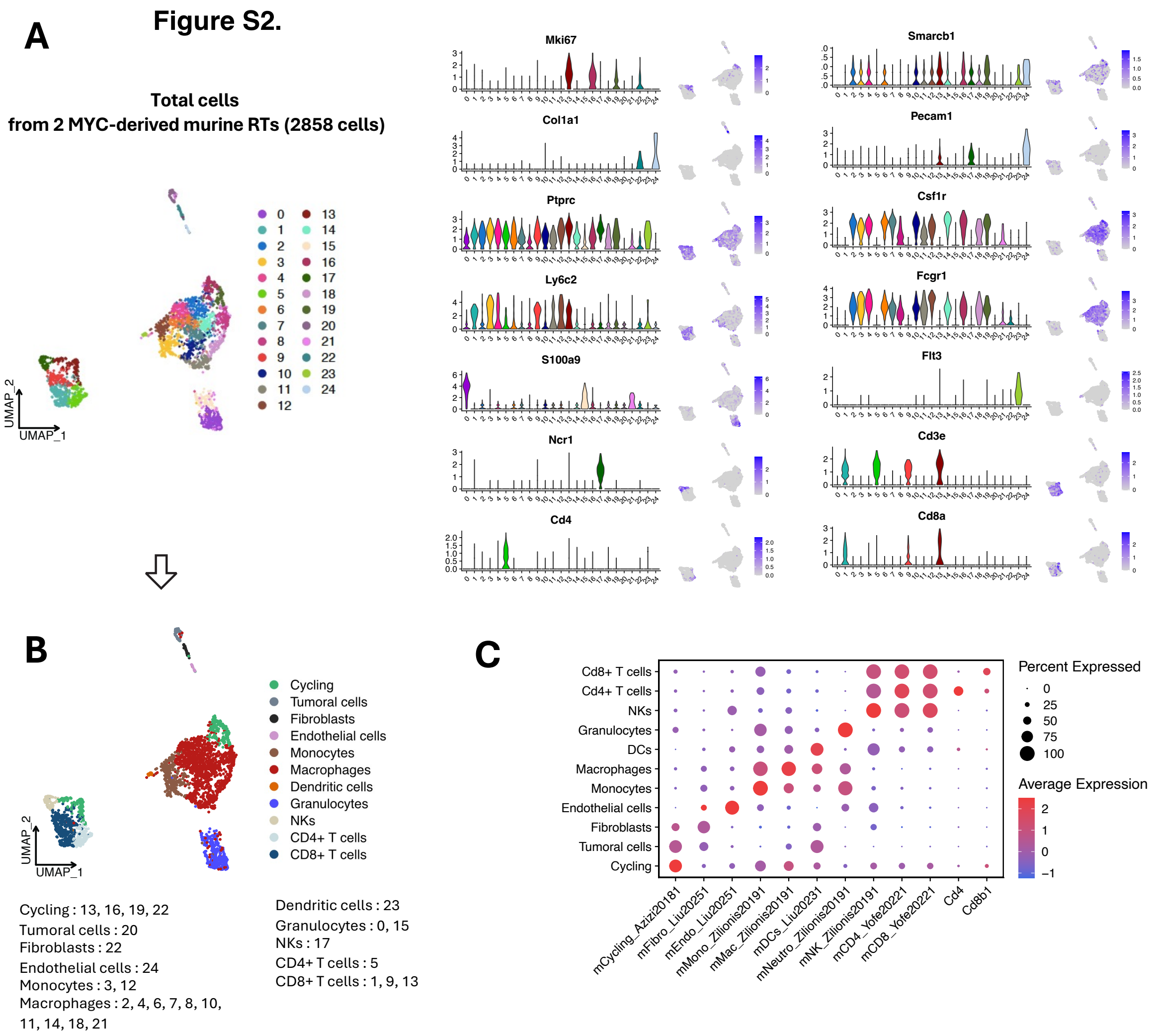
Figure S2.
A
Total cells
from 2 MYC-derived murine RTs (2858 cells)
UMAP_1
UMAP_2
Mki67
Smarcb1
Col1a1
Pecam1
Ptprc
Csf1r
Ly6c2
Fcgr1
S100a9
Flt3
Ncr1
Cd3e
Cd4
Cd8a
B
Cycling
Tumoral cells
Fibroblasts
Endothelial cells
Monocytes
Macrophages
Dendritic cells
Granulocytes
NKs
CD4+ T cells
CD8+ T cells
Cycling : 13, 16, 19, 22
Tumoral cells : 20
Fibroblasts : 22
Endothelial cells : 24
Monocytes : 3, 12
Macrophages : 2, 4, 6, 7, 8, 10, 11, 14, 18, 21
Dendritic cells : 23
Granulocytes : 0, 15
NKs : 17
CD4+ T cells : 5
CD8+ T cells : 1, 9, 13
C
Cd8+ T cells
Cd4+ T cells
NKs
Granulocytes
DCs
Macrophages
Monocytes
Endothelial cells
Fibroblasts
Tumoral cells
Cycling
mCycling_Azizi20181
mFibro_Liu20251
mEndo_Liu20251
mMono_Zilionis20191
mMac_Zilionis20191
mDCs_Liu20231
mNeutro_Zilionis20191
mNK_Zilionis20191
mCD4_Yofe20221
mCD8_Yofe20221
Cd4
Cd8b1
Percent Expressed
0
25
50
75
100
Average Expression
2
1
0
−1

**Fig. S2 (related to Fig. 2). Transcriptomic profile of the TIGIT/PD-1 axis in the RT mouse model supports the pre-clinical evaluation of dual immune checkpoint blockade.**

(A-C) Workflow for annotating cell populations from single-cell RNA-seq of total cells from two syngeneic RT tumor samples.

(A) UMAP from integrated total cells from two syngeneic RT tumor samples (2858 cells) at resolution 2.6 before annotation. Violin and feature plots showing the expression pattern of representative genes across clusters.

(B) Merging of similar clusters into consolidated and annotated cell populations based on shared gene expression profiles.

Clusters 13, 16, 19, 22 were merged into “Cycling”. Cluster 20 was renamed “Tumoral cells”. Cluster 22 was renamed “Fibroblasts”. Cluster 24 was renamed “Endothelial cells”. Clusters 3 and 12 were merged into “Monocytes”. Clusters “2, 4, 6, 7, 8, 10, 11, 14, 18, 21” were merged into “Macrophages”. Cluster 23 was renamed “Dendritic cells”. Clusters 0, 15 were merged into “Granulocytes”. Cluster 17 was renamed “NKs”. Cluster 5 was renamed “$CD4^+$ T cells”. Clusters 1, 9, 13 were merged into “$CD8^+$ T cells”.

(C) Dot plot showing the average gene expression per annotated cluster to validate cluster annotation.

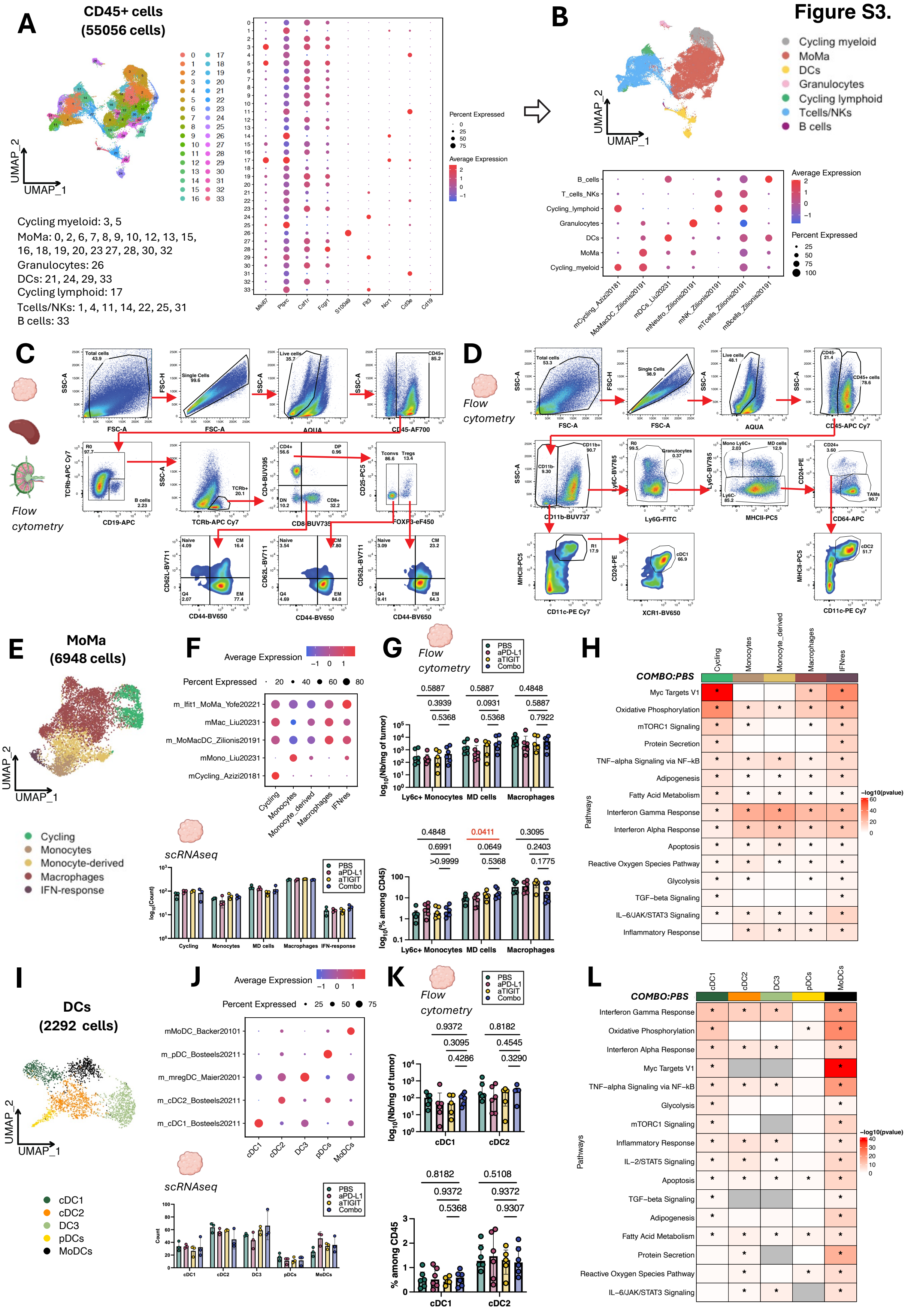
Figure S3.
A
CD45+ cells
(55056 cells)
UMAP_2
UMAP_1
Cycling myeloid: 3, 5
MoMa: 0, 2, 6, 7, 8, 9, 10, 12, 13, 15, 16, 18, 19, 20, 23 27, 28, 30, 32
Granulocytes: 26
DCs: 21, 24, 29, 33
Cycling lymphoid: 17
Tcells/NKs: 1, 4, 11, 14, 22, 25, 31
B cells: 33
Percent Expressed
Average Expression
Mki67
Ptprc
Csf1r
Fcgr1
S100a8
Flt3
Ncr1
Cd3e
Cd19
B
Cycling myeloid
MoMa
DCs
Granulocytes
Cycling lymphoid
Tcells/NKs
B cells
B_cells
T_cells_NKs
Cycling_lymphoid
Granulocytes
DCs
MoMa
Cycling_myeloid
mCycling_Azizi20181
MoMacDC_Zilionis20191
mDCs_Liu20231
mNeutro_Zilionis20191
mNK_Zilionis20191
mTcells_Zilionis20191
mBcells_Zilionis20191
C
Flow cytometry
Total cells
Single Cells
Live cells
CD45+
FSC-A
AQUA
CD45-AF700
CD19-APC
TCRb-APC Cy7
CD8-BUV735
FOXP3-eF450
CD44-BV650
CD62L-BV711
D
Flow cytometry
CD45-APC Cy7
CD11b-BUV737
Ly6G-FITC
MHCII-PC5
CD64-APC
CD11c-PE Cy7
XCR1-BV650
E
MoMa
(6948 cells)
Cycling
Monocytes
Monocyte-derived
Macrophages
IFN-response
F
Average Expression
Percent Expressed
m_Ifit1_MoMa_Yofe20221
mMac_Liu20231
m_MoMacDC_Zilionis20191
mMono_Liu20231
mCycling_Azizi20181
scRNAseq
PBS
aPD-L1
aTIGIT
Combo
G
Flow cytometry
Ly6c+ Monocytes
MD cells
Macrophages
H
COMBO:PBS
Myc Targets V1
Oxidative Phosphorylation
mTORC1 Signaling
Protein Secretion
TNF−alpha Signaling via NF−kB
Adipogenesis
Fatty Acid Metabolism
Interferon Gamma Response
Interferon Alpha Response
Apoptosis
Reactive Oxygen Species Pathway
Glycolysis
TGF−beta Signaling
IL−6/JAK/STAT3 Signaling
Inflammatory Response
Pathways
−log10(pvalue)
I
DCs
(2292 cells)
cDC1
cDC2
DC3
pDCs
MoDCs
J
mMoDC_Backer20101
m_pDC_Bosteels20211
m_mregDC_Maier20201
m_cDC2_Bosteels20211
m_cDC1_Bosteels20211
scRNAseq
K
Flow cytometry
% among CD45
L
COMBO:PBS
Interferon Gamma Response
Oxidative Phosphorylation
Interferon Alpha Response
Myc Targets V1
TNF−alpha Signaling via NF−kB
Glycolysis
mTORC1 Signaling
Inflammatory Response
IL−2/STAT5 Signaling
Apoptosis
TGF−beta Signaling
Adipogenesis
Fatty Acid Metabolism
Protein Secretion
Reactive Oxygen Species Pathway
IL−6/JAK/STAT3 Signaling

**Fig. S3 (related to Fig. 3). TIGIT/PD-L1 dual blockade enhances tumor rejection and survival in RT mouse model.**

(A-B) Workflow for annotating cell populations from single-cell RNA-seq of RT mouse tumors treated with PBS, anti-TIGIT, anti-PD-L1, or combined therapy (N=3 per group).

(A) Left: UMAP from integrated sorted $CD45^+$ cells from 12 RT murine tumor samples (55056 cells) at resolution 0.9 before annotation. Right: Dot plot showing the expression pattern of representative genes across clusters.

(B) Merging of similar clusters into consolidated and annotated cell populations based on shared gene expression profiles.

Clusters 3, 5 were merged into “Cycling myeloid”. Clusters 0, 2, 6, 7, 8, 9, 10, 12, 13, 15, 16, 18, 19, 20, 23, 27, 28, 30, 32 were merged into “MoMa”. Cluster 26 was renamed “Granulocytes”. Clusters 21, 24, 29, 33 were merged into “DCs”. Cluster 17 was renamed “Cycling lymphoid”. Clusters 1, 4, 11, 22, 25, 31 were merged into “Tcells/NKs”. Cluster 33 was renamed “B cells”.

(C-D) Gating strategies of flow cytometry analysis of the immune cells *in vivo* in n = 6-8 mice per treatment condition, treated with PBS, anti-TIGIT, anti-PD-L1, or combined therapy.

(C) Gating strategy using a lymphoid-focused panel for flow cytometry analysis of immune cells from tumors, spleens, and tumor-draining lymph nodes from mice treated with PBS, anti-TIGIT, anti-PD-L1, or combined therapy.

(D) Gating strategy using a myeloid-focused panel for flow cytometry analysis of immune cells from tumors from mice treated with PBS, anti-TIGIT, anti-PD-L1, or combined therapy.

(E, F, H) Single-cell RNA sequencing analysis of MoMa subsets isolated from Fig. 3B.

(E) UMAP visualization (N= 3 per treatment group; total cells=6948)

(F) Up panel: Dot plot showing the average gene expression per annotated cluster to validate cluster annotation; Down panel: quantification of MoMa subpopulations in single-cell MoMa subset. Bars represent mean values; error bars indicate the standard deviation (SD).

(G) Quantification (number/mg of tumor) and frequency (%) of MoMa subsets isolated from murine tumors (N=6) by flow cytometry across treatment groups. Bars represent the median cell count per group; error bars indicate the interquartile range (IQR).
(H) Pathway enrichment analysis comparing Combo- and PBS-treated tumors.
(I, J, L) Single-cell RNA sequencing analysis of DC subsets isolated from Fig. 3B.
(I) UMAP visualization (N= 3 per treatment group; total cells=2292)
(J) Up panel: Dot plot showing the average gene expression per annotated cluster to validate cluster annotation; Down panel: quantification of DC subpopulations in single-cell DC subset. Bars represent mean values; error bars indicate the standard deviation (SD).

(K) Quantification (number/mg of tumor) and frequency (%) of MoMa subsets isolated from murine tumors (N=6) by flow cytometry across treatment groups. Bars represent the median cell count per group; error bars indicate the interquartile range (IQR).
(L) Pathway enrichment analysis comparing Combo- and PBS-treated tumors.
Statistical significance was calculated using Mann-Whitney test, with significant results indicated in red in flow cytometry barplots and single-cell violin-plots and with a star (p-value < 0.05) in pathway enrichment heatmaps.

179 **Continuation of Supplementary Figure 3 (related to Figure 3).**

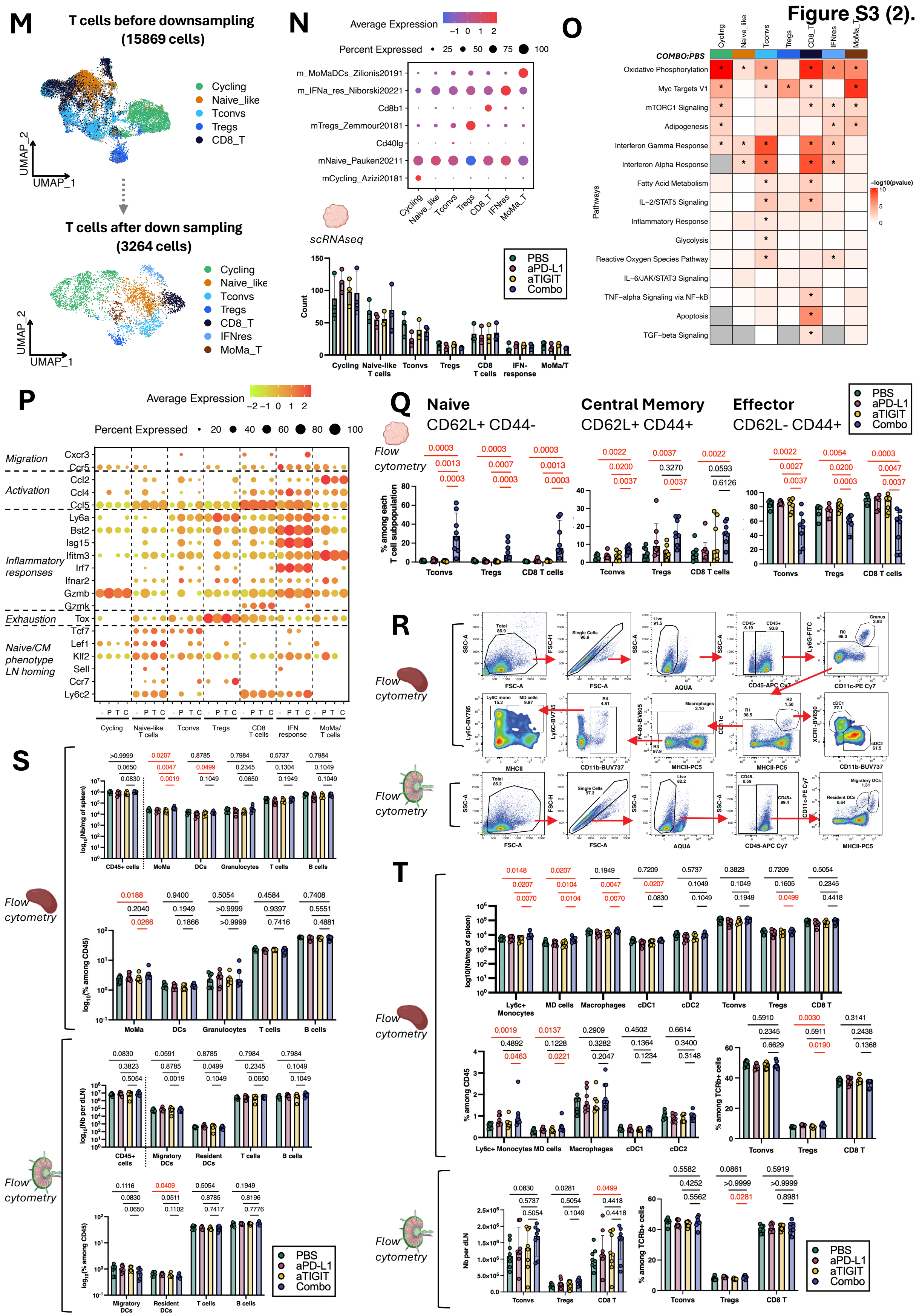


180

181

**Continuation of Fig. S3 (related to Fig. 3). Combination blockade of PD-L1 and TIGIT induces the formation of $PNAd^{+}$ HEV-like structures in the tumor microenvironment.**

(M-P) Single-cell RNA sequencing analysis of T cell subsets isolated from Fig. 3B.

(M) UMAP visualization from integrated T cells before down sampling (15869 cells) and after down sampling (3264 cells). N= 3 per treatment group.

(N) Up: Dot plot showing the average gene expression per annotated cluster to validate cluster annotation. Down: Quantification of T cell subpopulations in the single-cell RNAseq T dataset. Bars represent mean values; error bars indicate the standard deviation (SD).

(O) Pathway enrichment analysis comparing Combo- and PBS-treated tumors.

(P) Dot plots of selected gene expression profiles per treatment (- = PBS; P = anti-PD-L1; T = anti-TIGIT; C = COMBO) and per T cell subpopulations indicated on last row. Heatmap shows SCT-normalized expression per gene and cell. Dot size indicates the proportion of cells expressing the gene of interest within each cell type, and color indicates the level of expression from low (light green) to high (red). Only genes expressed in more than 20% of cells are shown.

(Q-T) Flow cytometry analysis of T cell subpopulations (Tconvs, Tregs, $CD8^{+}$ T cells) isolated from murine tumors and of major immune populations in spleen and tumor-draining lymph nodes (tdLNs). Bars represent the median cell count per group; error bars indicate the interquartile range (IQR). Statistical significance was calculated using Mann-Whitney test, with significant results indicated in red in flow cytometry plots.

(Q) Frequency (%) of Naive ($CD62L^{+}$ $CD44^{-}$), Central Memory ($CD62L^{+}$, $CD44^{+}$), Effector ($CD62L^{-}$ $CD44^{+}$) phenotype in Tconvs, Tregs and $CD8^{+}$ T cells.

(R) Gating strategy using a myeloid-focused panel for flow cytometry analysis of immune cells from spleens and tdLNs from mice treated with PBS, anti-TIGIT, anti-PD-L1, or combined therapy.

(S) Quantification (number/mg of spleen or tdLNs) and frequency (%) of major immune populations isolated from murine spleens and tdLNs by flow cytometry across treatment groups (N=8).

(T) Quantification (number/spleen or tdLNs), frequency of MoMa and DC subsets among $CD45^+$ cells (%) and frequency among $TCRb^+$ cells (%) of T cell subsets isolated from murine spleen and tdLNs by flow cytometry across treatment groups (N=8).

Statistical significance was calculated using two-tailed Mann-Whitney test, with significant results indicated in red in flow cytometry barplots and single-cell violin-plots and with a star ($p$-value $< 0.05$) in pathway enrichment heatmaps.

**Figure S4.**

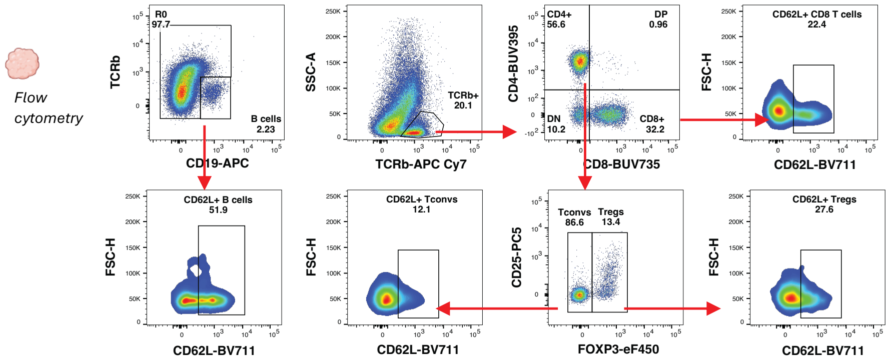


**Fig. S4 (related to Fig. 4). Gating strategy to visualize CD62L$^+$ B and T cells**

Gating strategy using a lymphoid-focused panel for flow cytometry analysis of immune cells from tumors from mice treated with PBS, anti-TIGIT, anti-PD-L1, or combined therapy.

### TABLES

#### Table S1. List of antibodies used in this study.

| Antigen | Fluorophore | Dilution | Clone | Catalog # | Company |
|---|---|---|---|---|---|
| NKp46 | FITC | 200 | 29A1.4 | 11-3351-82 | Thermo Fisher Scientific |
| TCF-1 | AF488 | 100 | C63D9 | 6444S | Cell Signaling Technology |
| CD39 | PeCy7 | 100 | 24DMS1 | 50-112-8745 | Thermo Fisher Scientific |
| CD25 | PC5 | 800 | PC61.5 | 15-0251-82 | Thermo Fisher Scientific |
| TCRb | APC Cy7 | 200 | H57-597 | 109220 | BioLegend |
| CD45.2 | AF700 | 200 | 104 | 109822 | BioLegend |
| CD19 | APC | 400 | 6D5 | 115512 | BioLegend |
| PD1 | BV785 | 100 | 29F.1A12 | 135225 | BioLegend |
| CD62L | BV711 | 200 | MEL-14 | 740660 | BD Biosciences |
| CD44 | BV650 | 400 | IM7 | 103049 | BioLegend |
| FoxP3 | eF450 | 100 | FJK-16s | 48-5773-82 | Thermo Fisher Scientific |
| CD4 | BUV395 | 200 | GK1.5 | 563790 | BD Biosciences |
| CD8 | BUV737 | 200 | 53-6.7 | 564297 | BD Biosciences |
| CD163 | Percp-E710 | 200 | TNKUPJ | 46-1631-82 | Thermo Fisher Scientific |
| Ly6G | FITC | 800 | 1A8 | 551460 | BD Biosciences |
| CD11c | PECy7 | 800 | N418 | 25-0114-82 | Thermo Fisher Scientific |
| MHCII (I-A I-E) | PC5 | 400 | M5/114.15.2 | 107612 | BioLegend |
| iNOs | PE-eF610 | 100 | CXNFT | 61-5920-82 | Thermo Fisher Scientific |
| CD24 | PE | 800 | M1/69 | 12-0242-81 | Thermo Fisher Scientific |
| CD45.2 | APC Cy7 | 100 | 104 | 109824 | BioLegend |
| Arg1 | Alexa 700 | 100 | A1exF5 | 56-3697-82 | Thermo Fisher Scientific |
| CD64 | APC | 100 | X54-5/7.1 | 139306 | BioLegend |
| CD11b | BUV737 | 800 | M1/70 | 612800 | BD Biosciences |
| Ly6C | BV785 | 800 | HK1.4 | 128041 | BioLegend |
| XCR1 | BV650 | 200 | ZET | 148220 | BioLegend |
| F4/80 | BV605 | 200 | BM8 | 123133 | BioLegend |
| CCR2 | BV421 | 200 | SA203G11 | 150605 | BioLegend |
| B220 | Purified | 400 | RA3-6B2 | 553086 | BD Biosciences |
| CD3 | Purified | RTU | Polyclonal | GA5036 | Agilent |
| CD11c | Purified | 200 | D1V9Y | 97585 | Cell Signalling |
| PNAd | Purified | 500 | MECA-79 | 553863 | BD Biosciences |
| TCF1/TCF7 | Purified | 100 | C63D9 | 2203 | Cell Signalling |
| CD31 | Purified | 100 | Polyclonal | ab28364 | Abcam |

234
